# Predicting transdermal fentanyl delivery using mechanistic simulations for tailored therapy

**DOI:** 10.1101/2020.06.16.154195

**Authors:** Thijs Defraeye, Flora Bahrami, Lu Ding, Riccardo Innocenti Malini, Alexandre Terrier, René M. Rossi

## Abstract

Transdermal drug delivery is a key technology for administering drugs. However, most devices are “one-size-fits-all”, even though drug diffusion through the skin varies significantly from person-to-person. For next-generation devices, personalization for optimal drug release would benefit from an augmented insight into the drug release and percutaneous uptake kinetics. Our objective was to quantify the changes in transdermal fentanyl uptake with regards to the patient’s age and the anatomical location where the patch was placed. We also explored to which extent the drug flux from the patch could be altered by miniaturizing the contact surface area of the patch reservoir with the skin. To this end, we used validated mechanistic modeling of fentanyl diffusion, storage, and partitioning in the epidermis to quantify drug release from the patch and the uptake within the skin. A superior spatiotemporal resolution compared to experimental methods enabled *in-silico* identification of peak concentrations and fluxes, and the amount of stored drug and bioavailability. The patients’ drug uptake showed a 36% difference between different anatomical locations after 72 h, but there was a strong interpatient variability. With aging, the drug uptake from the transdermal patch became slower and less potent. A 70-year-old patient received 26% less drug over the 72-h application period, compared to an 18-year-old patient. Additionally, a novel concept of using micron-sized drug reservoirs was explored *in silico*. These reservoirs induced a much higher local flux (µg cm^-2^ h^-1^) than conventional patches. Up to a 200-fold increase in the drug flux was obtained from these small reservoirs. This effect was mainly caused by transverse diffusion in the stratum corneum, which is not relevant for much larger conventional patches. These micron-sized drug reservoirs open new ways to individualize reservoir design and thus transdermal therapy. Such computer-aided engineering tools also have great potential for *in-silico* design and precise control of drug delivery systems. Here, the validated mechanistic models can serve as a key building block for developing digital twins for transdermal drug delivery systems.

## 1 INTRODUCTION

Transdermal drug delivery systems (TDDS) are used to deliver non-invasively moderately lipophilic, low-molecular-weight drugs to patients. Delivery through the skin, the largest human organ, enables controlled drug administration to achieve steady blood plasma concentrations without a distinct concentration peak [1]. Via the skin, drugs that have low bioavailability via oral administration, due to a high first-pass effect, can be effectively delivered. The ideal molecule for transdermal drug delivery has a molecular weight below 500 Da and a log partition coefficient (octanol-water) of 1 to 3 [2], [3], so balanced lipophilicity. These molecules can cross the lipophilic stratum corneum but also diffuse through the hydrophilic viable epidermis and dermis, into the aqueous systemic circulation [2]. Commercial TDDS have been developed for fentanyl, nitroglycerin, estradiol, nicotine, and testosterone, amongst others. In this multi-billion US dollar market [4], [5], transdermal fentanyl patches currently provide the highest product sales, as a popular solution for around-the-clock opioid analgesia [2].

A key hurdle in these TDDS is that the pathway for delivery —the human skin— is very patient-specific. Absorption kinetics depends on the individual patient’s skin composition and the hygro-thermo-mechanical properties of each of its sublayers. Additionally, the patient’s metabolism, lifestyle, and bio-environment play a role. Studies have quantified the variability in drug uptake with body location [6]–[8], age [8]–[10], gender [8], ethnicity [11], skin hydration and disease state [12] as well as among patients within the same subject category [6]. The interpatient variability in the plasma concentration is caused by the complex combination of the TDDS release, transdermal drug absorption, circulation within the body and drug elimination. Any variation in these steps leads to inter- or intra-individual variability in the effect of the drug. The identified variability is dependent on the specific drug molecule and the composition and size of the target group [13]. The interpatient variability was thereby found to be larger or smaller than with oral delivery, depending on the drug [13]. This patient-induced variability makes precise transdermal dosage challenging. Clinical repercussions are that the therapeutic drug level in the blood is not always reached with certain patients or overdosing occurs when blood concentrations are out of the therapeutic range. Typical side effects are ineffective pain relief, skin irritation, respiratory depression, apnea, or, in some extreme cases, death [14], [15].

Instead of “one-size-fits-all”, future TDDS and corresponding control strategies are designed to provide drugs for each patient at the correct rate for a specific body location [16], [17]. Defining a patient-specific therapeutic window was already proposed for certain drugs, including fentanyl [18], since also the minimum effective concentration differed significantly between patients. However, personalizing transdermal drug delivery requires a clear insight into release and uptake kinetics and the associated biophysical processes that drive the uptake. Current transdermal drug delivery experiments have a rather low spatiotemporal resolution, large inter-sample variability, and are often performed for infinite drug reservoirs [19]. A typical example is monitoring the cumulative drug uptake in Franz diffusion cells by analyzing samples with high-performance liquid chromatography (HPLC) at time intervals of several hours. Another example is obtaining steady-state concentration-depth profiles over the skin via tape stripping [6], [20], [21]. In several experiments, the skin samples are exposed to water reservoirs for a prolonged time. Water acts as a penetration enhancer for the skin and is often used in clinical dressings to increase drug diffusion [22]. Therefore, the diffusion coefficients determined via Franz diffusion cells will represent an upper limit.

Mathematical modeling is an alternative method to gain complementary insights into the transport processes and to design and optimize next-generation TDDS *in silico*. The following methods have been used in the literature:

1. Mechanistic models that solve partial differential equations, for example at the macroscale, mesoscale [23] and even cellular level (microscale; Figure 1 a, b, c) with finite elements [24].
2. Molecular dynamics at the subcellular level (e.g., lipid layers) to obtain transport properties and thermodynamic values of the system (e.g., partition coefficients) [25] that could then be used in a multiscale approach [26], [27] (Figure 1d). Additionally, alternative modeling strategies have been proposed that account for interactions at the molecular level [28] to calculate the skin permeability.
3. Compartmental models, which solve ordinary differential equations and consider each skin layer as a well-mixed compartment [19], [29], [30]. Usually, there is no discretization over the skin layer, although sometimes the skin layer is subdivided into a few compartments.

Mechanistic models are of particular interest, compared to analytical solutions of drug uptake in the skin. Analytical models of the diffusion process are only valid for simple boundary conditions, which typically do not change in time or space and are derived for simple geometries [32]. This impedes calculating, for example, a transdermal therapy that changes over time. Also, analytical solutions typically only calculate diffusion, without the presence of any physical binding or metabolization of the drug molecule in the skin. Mechanistic models provide drug concentrations and flow inside the skin and drug reservoir at each point in space and time in three dimensions, to the specific degree of detail that is dependent on the used method. Finite element, finite volume, or finite difference methods are typically used. These methods enable researchers to quantify, among other things, the time lag in drug release and subsequent uptake in the blood, local peak concentrations or fluxes, high-resolution concentration-depth profiles [33], the depletion state of the drug reservoir, and the remaining drug amount stored in the skin. Mechanistic models also enable one to explore a large parametric space of process variables. Considerable work has been performed on mechanistic modeling for TDD (Table 1). These studies, however, do not explicitly focus on intra- or interpatient variability, a key step towards tailoring TDDS for individual patients or categories of patients, e.g., different age groups.

**Table 1.**
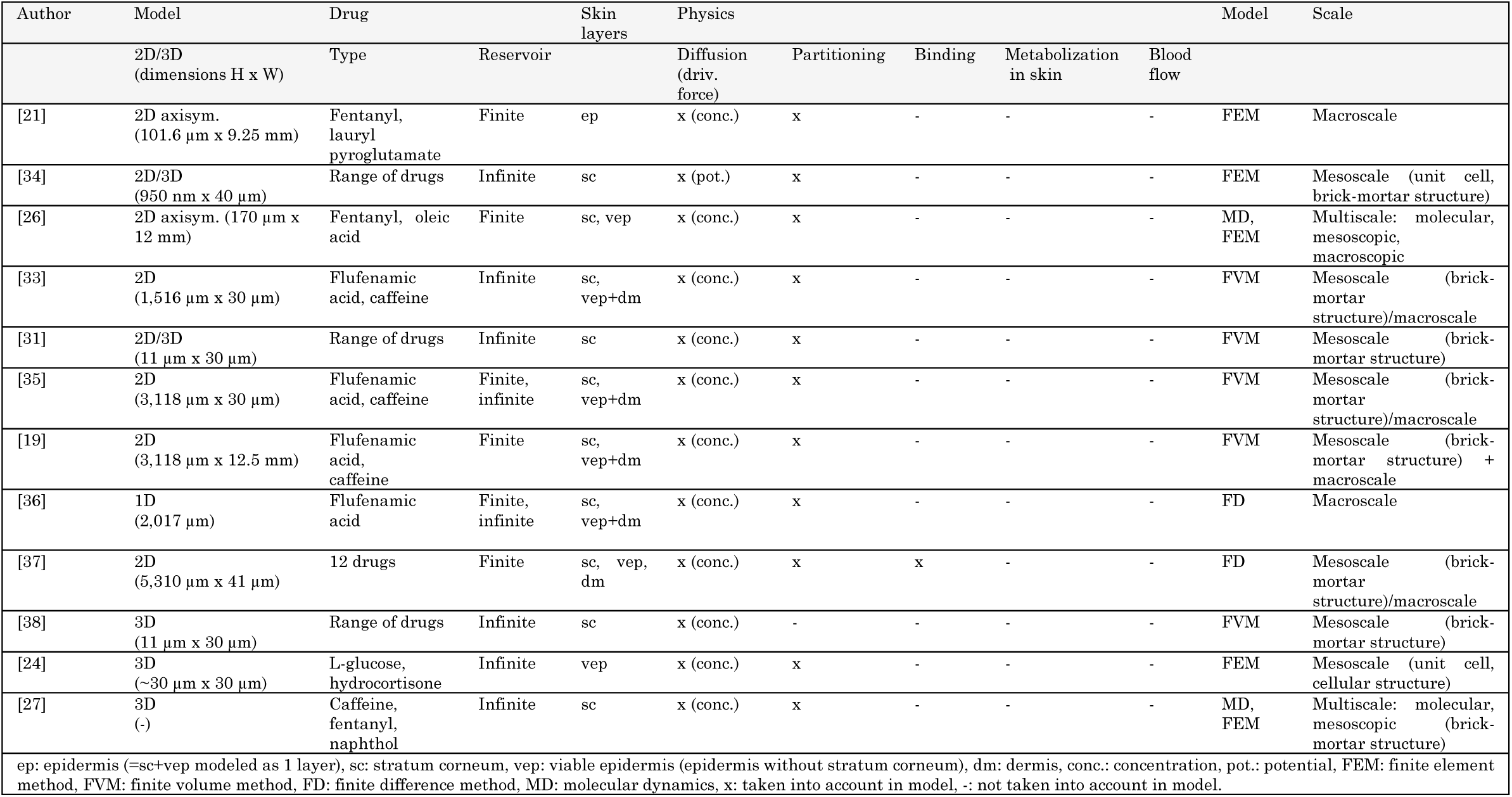
Non-exhaustive overview of mechanistic modeling work of transdermal drug delivery over the past decade (ordered chronologically).

**Figure 1.**
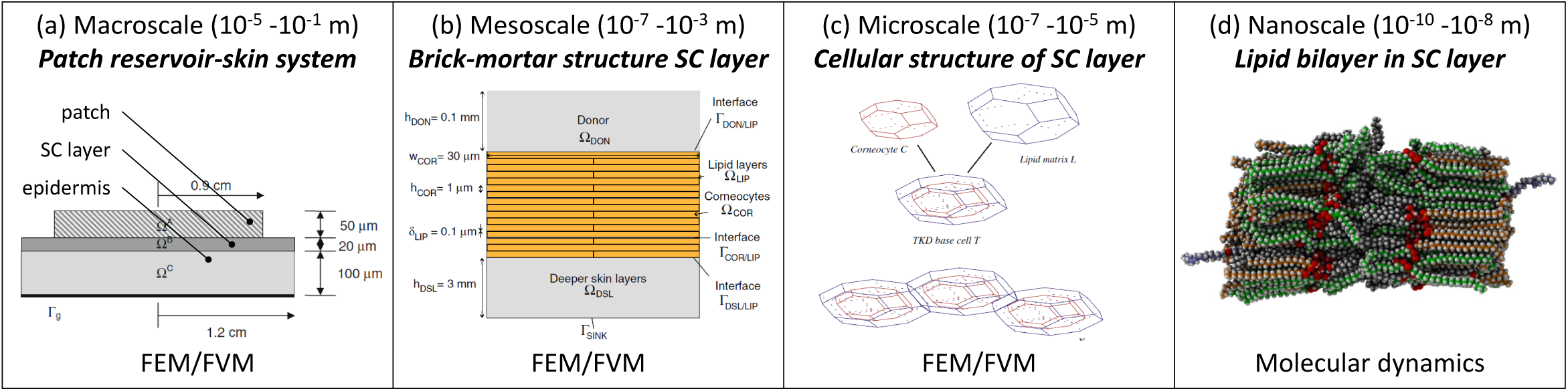
Overview of existing mechanistic models (adapted from (a) [26], (b) [19], (c) [31]).

In this study, our objective was to quantify the changes in transdermal fentanyl uptake with the patient’s age and the anatomical location where the patch was placed. We also explored how the drug flux from the patch could be altered as a function of the contact surface area of the patch reservoir with the skin. We applied mechanistic modeling to quantify *in-silico* differences between the scenarios. A finite-element model for transdermal drug delivery was developed in line with the current state-of-the-art. This mechanistic model was validated, and the sensitivity to the model parameters evaluated. With this model, we took three steps beyond the current state-of-the-art. First, we answered how much more fentanyl is taken up transdermally by a specific patient who applies the same patch now and 50 years in the future (due to skin aging). Second, we quantified how much more effective, or harmful, a transdermal patch is when placed at a different anatomical location. Third, we evaluated how the drug flux can be enhanced by miniaturizing the drug reservoir size, a process that takes advantage of transverse diffusion in the stratum corneum layer.

## 2 MATERIALS AND METHODS

### 2.1 Continuum model for transdermal fentanyl delivery

#### 2.1.1 Computational system configuration

A mechanistic continuum model was built to simulate fentanyl release from a transdermal patch (reservoir) and subsequent uptake through healthy human skin. This study targets transdermal delivery of fentanyl, one of the most common drugs delivered transdermally [2]. This synthetic opioid is approximately 100 times more potent than morphine [39]. Fentanyl transdermal patches are used for patients with severe chronic pain, like cancer patients during treatment or at their end-stage [40]. Fentanyl has the appropriate lipophilicity and is sufficiently small to readily penetrate through the skin barrier and then reach the blood circulation. Therefore, conventional, first-generation transdermal patches can be used, where the uptake process through the skin barrier is diffusion-driven [41]. The model and simulation were built and executed according to best practice guidelines in modeling for medical device design [42], [43].

The geometrical setup, along with the boundary conditions, is depicted in Figure 2. The system configuration includes a square-shaped drug reservoir, which contains a finite amount of fentanyl, and the outer part of the human skin, namely the epidermis. TDDS are designed to deliver drugs at a nearly constant rate, similar to other controlled-release dosage systems [1], [44]. The fentanyl patches, therefore, are commercially labeled with the targeted drug release rate, typically 12-100 µg h^-1^ (Table 2, [45]) over the 72-h application period. Higher delivery rates are obtained by simply increasing the contact surface area of the patch (range between 4.2-42 cm^2^, Table 2). Conventional transdermal fentanyl therapy consists of estimating empirically the initial dose (so patch size) for the patient, applying the patch transdermally, and replacing it every 72 hours [40]. Patches are not allowed to be resized manually ad posterior to change the dose, as it may lead to a fast initial release of fentanyl [69].

**Table 2.** Delivery rate, active surface contact area, and drug content of selected, commercially-available transdermal patches. The flux was deduced from the labeled delivery rate and active area of the patch through which the drug is released.

| Commercial product | Targeted delivery rate ($G_{pt,rel}$) $\mu\text{g h}^{-1}$ | Active area ($A_{pt}$) $\text{cm}^2$ | Drug content mg | Drug flux ($G_{pt,rel}/A_{pt}$) ($\mu\text{g cm}^{-2}\ \text{h}^{-1}$) | Reference |
| --- | --- | --- | --- | --- | --- |
| Durogesic® | 12-100 | 5.3-42 | 2.1-16.8 | 2.3-2.4 | [2], [57] |
| Matrifen® | 12-100 | 4.2-33.6 | 1.4-11 | 2.9-3.0 | [2], [57] |
| Fentalis® | 25-100 | 10-40 | 2.5-10 | 2.5 | [2], [57] |
| Fentadolon® | 25 | 15 | - | 1.7 | [2] |

**Figure 2.**
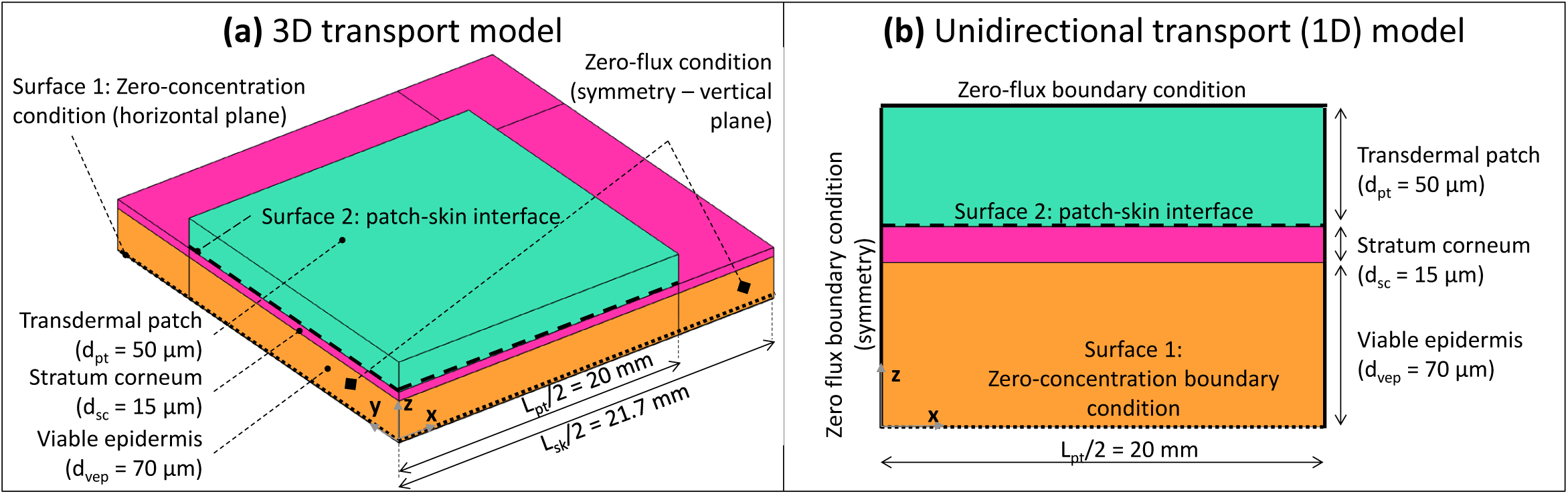
3D and 1D geometrical models of square-shaped drug reservoir and skin (not to scale; for 3D model, only one-fourth of the system was modeled due to the symmetry).

For the drug reservoir, a patch length *L*_*pt*_ of 40 mm was chosen. This value implies an active area of 16 cm^2^, lying within the range reported for commercial transdermal patches for fentanyl (4.2-42 cm^2^, Table 2, [2]). The thickness of the patch (*d*_*pt*_) was chosen to be 50 μm. Thus the volume of the patch reservoir was 80 mm^3^. These dimensions lead to a realistic initial drug content in the reservoir (mg), based on the initial concentration, as shown in section 2.1.4. The skin’s epidermis (ep) was composed of two layers: the stratum corneum (sc) and the viable epidermis (vep). The reason for this model configuration is that the lipophilic stratum corneum — a brick-mortar structure of lipid bilayers and corneocytes — has markedly different transport properties compared to the hydrophilic viable epidermis [46]. The stratum corneum exhibits the primary mechanical barrier function for drug delivery through the skin, together with tight junctions [46]–[49], which are located inside the viable epidermis. The impact of tight junctions on the drug diffusion was not modeled separately but rather lumped into the transport properties of the epidermis. Tight junctions are rarely included explicitly in existing mechanistic models [47].

The dermis could also play a role in drug transport and drug retention during transdermal delivery [50]. Previous works modeled the dermis (1) as an additional depot in which drugs can be stored [51]; (2) as an additional layer in which only diffusion and no drug removal by the blood flow was modeled [52]; (3) by adding a sink condition to the system [53]; (4) by modeling simplified capillary loops in the dermis [54]. An even more realistic model of dermis would require modeling the patient’s blood flow since capillaries and vessels are present in the papillary and reticular dermis, respectively. Modeling blood flow would also require including the associated biochemical processes in the capillaries, an undertaking that was beyond this study’s scope. Therefore, the dermis was not included in the computational domain in the present study.

Two computational domains were constructed: (1) a one-dimensional (1D) model, where the drug reservoir was as wide as the skin, so this simulated unidirectional drug transport (Figure 2b). This case is representative of Franz diffusion cell experiments [6], [21], [55] and has a low computational cost; (2) a three-dimensional (3D) model where the skin was wider than the drug reservoir to capture possible transverse diffusion of the drug (x and y directions in Figure 2a), since the drug spreads laterally, the actual delivery surface area becomes larger than the patch surface area [56]. The skin was extended by 1.7 mm (= 20 x (d_sc_ + d_vep_)) on each side of the patch to avoid an impact of the transverse boundary on the simulated drug uptake kinetics at the interface between epidermis and dermis (surface 1 in Figure 2), as determined by a sensitivity analysis. Only one-fourth of the geometry was explicitly modeled by considering the symmetry in the system.

#### 2.1.2 Governing equations

Only the diffusion of fentanyl was solved, and isothermal conditions were assumed, namely, ambient body temperature. Neither water transport due to skin de-/rehydration, nor the resulting skin shrinkage or swelling was included. To derive the mass conservation equation, we started from the following equation, defined for each material *i* [kg m^-3^] to describe the drug concentration 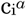 of substance *α*:

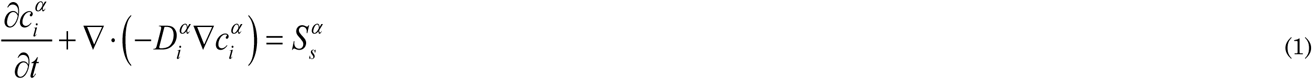

where 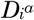 is the corresponding diffusion coefficient or diffusivity [m^2^ s^-1^], 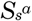 is a volumetric source term for substance α [kg m^-3^s^-1^], and *t* is the time [s]. No source term was included in this study since the contributions of the following processes could be neglected [23]:

– Clearance of the drug via the blood capillaries and vessels into the body. As only the epidermis was included in the model (section 2.1.1), in which no blood capillaries or larger blood vessels are present, this effect was not included.
– Metabolization of the drug molecule within the skin by chemical reactions that may lead to a conversion of the drug into other compounds. Fentanyl is mainly metabolized in the liver and also by the intestinal mucosa for oral delivery, but not within the skin [58], [59]. Therefore, metabolization within the skin is low and was not modeled.
– Adsorption of the drug molecule into the skin, and thus physical binding of the drug molecules. This process is different from the storage of unbound drug molecules in the tissue during transient drug uptake. The bound molecules could sometimes also unbind later and diffuse into the blood circulation. Adsorption of the drug in the skin is one of the processes that reduce the bioavailability of the drug. Bioavailability is the amount of drug administered through the skin that reaches the systemic circulation in an unchanged state. Bioavailability is a key pharmacokinetic characteristic that is determined by physical adsorption and chemical metabolization processes of the drug in the skin. The bioavailability of fentanyl transdermal delivery is very large (e.g., 92% in [60]). This means that most of the drug which diffuses from the patch will reach the blood circulation. By considering this fact, we can conclude that one of the processes reducing this bioavailability – adsorption of fentanyl molecules in the skin – is also rather limited. Therefore, as an approximation, no physical adsorption source term was modeled in the skin domain.

A key phenomenon during drug uptake in multi-layer assemblies, such as the skin, is drug partitioning [61]. Partitioning implies that when a drug α is brought into contact with two materials (A and B), the drug concentration in these two materials will equilibrate to different values, namely 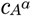 and 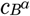. The ratio of these equilibrium concentrations is called the partition coefficient 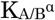.

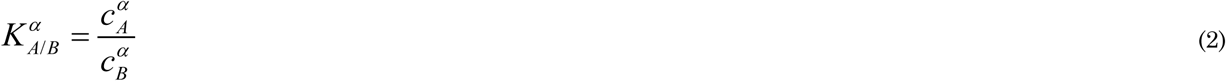

For drug partitioning in liquids, the octanol-water partition coefficient is often determined 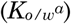, where values larger than 1 indicate drug lipophilicity and values smaller than one indicate drug hydrophilicity. The log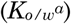 is also often reported, where positive/negative values indicate lipophilicity/hydrophilicity, respectively.

Partitioning leads to a discontinuity in the drug concentration at the interface between the two materials, for example, between the stratum corneum and the viable epidermis. This discontinuity can affect the numerical stability of the simulation. An elegant solution is to substitute the dependent variable drug concertation 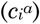 for another variable 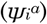 in Eq.(1) in the following way [23], [34]:

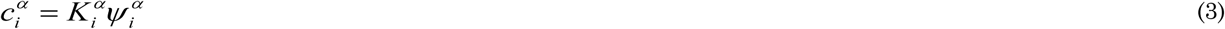

where 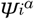 is termed in this study, the drug potential for every material *i* and is defined in [kg m^-3^]. 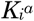 is termed the drug capacity of the drug in the material *i* [-]. Similar substitutions were made in other research fields [62]–[64]. This choice of dependent variable avoids numerical stability issues. This substitution is elaborated in Supplementary Material 1 and results in a single mass conservation equation instead of one for each material (Eq.(1)):

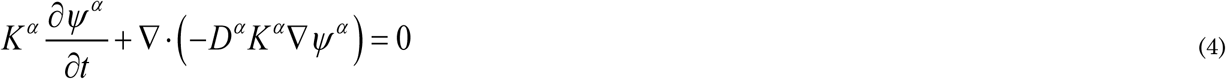

where 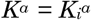 and 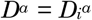 for each material *i*, so these parameters were defined separately for each material. The dependent variable *ψ*^*α*^ is continuous throughout all materials and over all the interfaces [23]. The partition coefficient can also be defined as the ratio of drug capacities by combining Eq.(2) and Eq.(3):

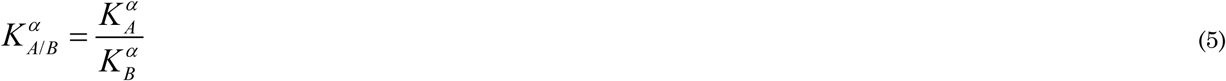

Note that at all interfaces between different materials, continuity of fluxes is inherently maintained:

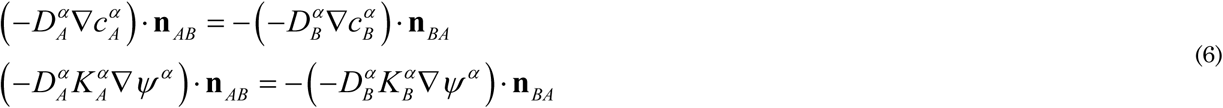

where **n** is the unit vector normal to the interface.

#### 2.1.3 Material properties and transport characteristics of skin and patch

Fentanyl is a synthetic opioid used as a pain medication. It has a low molecular weight (337 Da) and is moderately lipophilic with a log(*K*_*o/w*_) of 3 to 4 [2], [26], [58]. The material transport properties of the skin components and the drug reservoir are given in Table 3 for fentanyl for the different cases that were simulated (see section 2.2). Values were taken from the literature [21], [26]. The different material property data sets that were used (validation study versus parametric study) led to similar uptake kinetics (see Supplementary Material 3). The drug capacities are derived from the partition coefficients via Eq.(5), as the latter are typically available from measurements. To be able to do so, one drug capacity needed to be fixed, and therefore 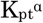 was set (arbitrarily) equal to one. Similar to most other simulation studies [26], [27], [35], the diffusion and partition coefficients were taken as constants and isotropic, meaning that they are independent of the drug concentration because more detailed data was not available.

**Table 3.**
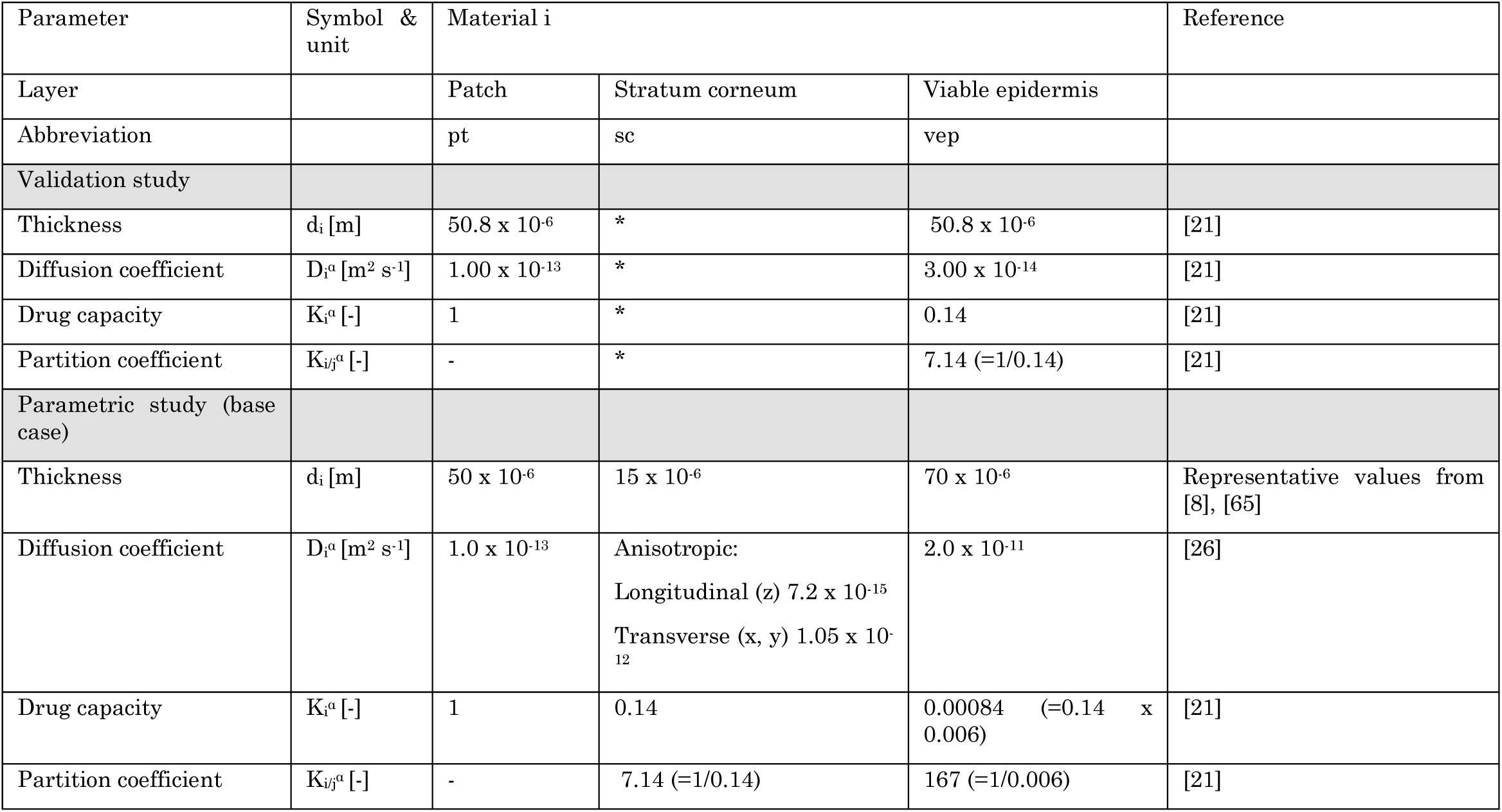
Material transport properties used in the model for different configurations for fentanyl. (*: not explicitly modeled but included in the epidermis)

#### 2.1.4 Boundary and initial conditions

The drug was assumed to be removed from the computational domain only via the interface with the dermis. In the dermis, it will be transported by diffusion and blood flow. To this end, a constant concentration (and potential), equal to zero, was imposed at the bottom of the epidermis (Figure 2), as in previous studies [21], [34]. This condition represents a Dirichlet boundary condition. This assumption is justified by the fact that the drug concentration in the dermis is very low because of a higher drug diffusion coefficient, drug partitioning between dermis and epidermis, and the fact that the blood flow extracts drugs. Zero-flux conditions were imposed at all vertical boundaries. At t=0, the skin was assumed to be drug-free. The initial drug concentration in the patch was set at 80 kg m^-3^, according to a previous study [21]. This dose implies a total initial amount of fentanyl in the reservoir m_pt,ini_ = 6.4 mg, which corresponds to amounts typically present in commercially available patches (Table 2). Complete contact between the patch and the skin was assumed, without any inclusion of air layers or discontinuities like hairs or skin roughness.

### 2.2 Spatial and temporal discretization

The finite element mesh was built based on a grid sensitivity analysis on the 1D and 3D models. Using Richardson extrapolation, the spatial discretization error on the total mass flux to the dermis was estimated to be 0.1% for both 1D and 3D models [66], [67]. The grid consisted of 120 quadrilateral finite elements (1D, elements with a size of about 1 µm) and 107,000 hexahedral finite elements (3D) for the base case. The grid was gradually refined towards the different material interfaces to enhance numerical accuracy and stability, because the largest gradients are found at such interfaces, particularly at the initial stage of the uptake process.

Starting from these initial conditions, the transient simulations calculated a drug uptake process that lasted 240 h (10 days), to capture the drug uptake history until depletion. These simulations applied adaptive time-stepping, with a maximal time step of 600 s (10 min). This time step was chosen to ensure high temporal resolution for the output data and was determined from a sensitivity analysis.

### 2.3 Alternative configurations

The base case (Table 3) mimicked the drug uptake in the skin, as released from a finite drug reservoir. Additional configurations were simulated with partially different geometries, process conditions, or reservoir/skin material properties:

– Validation of the drug release and uptake with experimental data from a previous study [21]. A detailed description of the experiments and the corresponding simulations is given in Supplementary Material 2 and Table 3.
– Sensitivity analysis to multiple model parameters (detailed in section 2.3.1)
– An infinite reservoir with a very high diffusion coefficient that cannot be depleted (detailed in section 2.3.2).
– The case where the patch was removed after 72 h, which was performed by changing the boundary conditions (detailed in section 2.3.2).
– Variability with respect to the anatomical location where the patch was placed on the patient’s body and the age of the patient (detailed in section 2.3.3).
– Reservoir contact surface area and size, in terms of its transverse dimensions (x,y in Figure 2; detailed in section 2.3.4).

Unless specified otherwise, 1D unidirectional models were used (Figure 2). To support this decision, the differences with a 3D model and the impact of the anisotropic transport properties between transverse and longitudinal directions were quantified for the evaluated patch widths in Supplementary Material 3 and section 3.1.

#### 2.3.1 Sensitivity to model parameters

The sensitivity of the drug uptake process to various model parameters was explored to identify the ones with the largest impact. To this end, the relative sensitivity *S*_*U, Xj*_ of a process quantity *U(t)* (e.g., drug flux) to a change in a model input parameter *X*_*j*_ over time was calculated using partial derivatives:

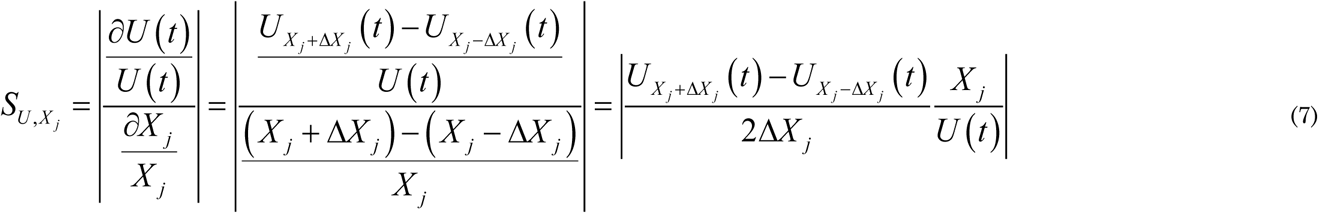

where *ΔX*_*j*_/*X*_*j*_ was set equal to a 1 % deviation from the nominal value of *X*_*j*_. The sensitivity to the following parameters (*X*_*j*_) was probed: 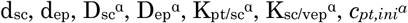. The following quantities, *U(t)*, were evaluated at specific time points (12, 24, 48 and 72 h): (1) the uptake flux across the skin into the dermis, so into the blood g_bl,up_ [kg m^-2^ s^-1^]; (2) the total amount of drugs taken up via the dermis by the blood flow m_bl,up_(t) [kg]. Such a sensitivity analysis is an essential step to designing drug delivery systems and therapies that achieve a constant drug delivery for the patient.

#### 2.3.2 Infinite reservoir and patch removal after 72 h

An infinite reservoir with a very high diffusion coefficient was simulated. As this reservoir cannot be depleted, the gradient over the skin remains constant once a steady state is reached. To this end, the boundary conditions were adjusted. For these simulations, the patch was removed in the system configuration and a constant concentration 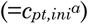 was imposed at surface 2 (Figure 2). To simulate the removal of the patch after 72 h, a zero-flux condition was imposed at surface 2, and only transport in the epidermis was solved.

#### 2.3.3 Patient-specific parameters

The patient-specific uptake of fentanyl through the skin epidermis was quantified in two steps. First, the intra-patient variability was analyzed, namely by investigating the effect of the anatomical application site on the human body of the transdermal drug delivery device. The different body sites that were tested and their corresponding stratum corneum and viable epidermis thickness are presented in Table 4. Simulations were performed with the average thickness (µ_i_), but also with thicknesses that corresponded to µ_i_ -/+ 2σ_i_, respectively, where σ_i_ is the standard deviation of material *i*. This range corresponds to the 2.5% and 97.5% quantiles of the sublayer thickness, and, thus, about 95% of the thickness range was covered.

**Table 4.** The thickness of stratum corneum and viable epidermis at different body sites - mean (standard deviation). The data were taken from [8], who evaluated 71 subjects (37 males, 34 females, age: 20-68 years, median: 47 years) and reported the data on each patient.

| Parameter | Stratum corneum ($\mu\text{m}$) | Viable epidermis ($\mu\text{m}$) | Total epidermis ($\mu\text{m}$) |
| --- | --- | --- | --- |
| Forearm dorsal | 18.3 (4.9) | 56.6 (11.5) | 74.9 |
| Shoulder | 11.0 (2.2) | 70.3 (13.6) | 81.3 |
| Buttock | 14.9 (3.4) | 81.5 (15.7) | 96.5 |
| All body sites | 14.8 (4.8) | 69.9 (17.0) | 83.7 |

Second, the interpatient variability on the drug uptake kinetics was evaluated, namely by assessing patients from different age categories. The stratum corneum thickness increases with age [10]. In addition to the thickness of the stratum corneum, which is the main barrier for transdermal drug delivery, also the composition of the skin changes, where the lipid content changes. These compositional changes, in turn, affect the diffusion coefficient and the partition coefficient. Previous studies suggest that with increasing age, the diffusion coefficient, and thus the permeability of the skin slightly decreases [10], [68]. This decrease in permeability of the skin does not significantly affect the permeation for lipophilic drugs [9]. Therefore, in this study the possible changes in the composition of the stratum corneum layer due to age and the resulting diffusion and partition coefficient were not investigated. Note that these effects would even strengthen the reduction of transdermal drug uptake with age. Only the effect of age on the stratum corneum thickness was included, so only the skin layer thicknesses were altered, while the other material properties were assumed to remain constant. Identifying the intra- and interpatient variability of the drug uptake kinetics will help to design drug delivery systems and therapy so that every patient can receive an optimal drug dose.

The following correlation was used to relate the stratum corneum thickness (d_sc_ [m]) and age (A [a], i.e., years) for the dorsal forearm, as reported previously [10] (40 female subjects, 10 in each of the following age categories: 18–30, 30–40, 40–55, and 55– 70 years):

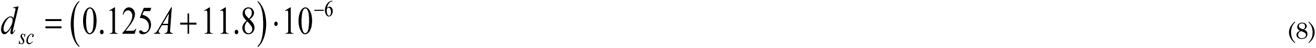

#### 2.3.4 Reservoir contact surface area

In addition to the 40 mm wide drug reservoir, other reservoir sizes were evaluated. The rationale for this decision was to quantify how the contact surface area (patch length or width *L*_*pt*_) of the patch affects the drug flux to the skin. The following patch widths were evaluated: 40 mm (base case), 4 mm, 400 µm, 40 µm, 4 µm. Since smaller reservoirs are depleted faster, simulations with an infinite reservoir (section 2.3.2) were also conducted.

### 2.4 Numerical implementation and simulation

The model was implemented in COMSOL Multiphysics® software (version 5.4, COMSOL AB, Stockholm, Sweden), a finite-element-based commercial software program. This software was verified by the code developers. Therefore, additional code verification was not performed by the authors. Transient diffusive drug transport (Eq.(4)) in the patch and skin during drug release and uptake was solved using the partial differential equations interface (coefficient form). The conservation equation was solved for the dependent variable *ψ*. Quadratic Lagrange elements were used with a fully-coupled direct solver, which relied on the MUltifrontal Massively Parallel sparse direct Solver (MUMPS) solver scheme. The tolerances for solver settings and convergence were determined by means of sensitivity analysis in such a way that a further increase in the tolerance did not alter the resulting solution.

### 2.5 Metrics

The simulated drug delivery process was analyzed quantitatively by calculating several metrics. With respect to the stored drug amount, we quantified the remaining (residual) amount contained in the patch as a function of time (m_pt,res_(t) [kg]), and the total amount of drugs stored (present) in the skin (viable epidermis and stratum corneum) as a function of time (m_ep,stor_(t) [kg]).

For the transported drug quantity, the drug amount released by the patch (m_pt,rel_(t) [kg]) was quantified as a function of time, which is the cumulative integrated flux over surface 2 (Figure 2) over time. The corresponding release flow rate (G_pt,rel_ [kg s^-1^]), namely the slope of the m_pt,rel_(t) curve, was also determined. Furthermore, the drug amount that was taken up by the dermis was also determined. Due to the large bioavailability of fentanyl, we assumed that the drugs that exit the computational system. Hence, the amount uptaken by the dermis equals the drugs taken up into the blood flow. Using the outgoing drug flow, the drug amount that was taken up by the blood flow, m_bl,up_(t) [kg], was determined as a function of time. It is the cumulative integrated flow over surface 1 (Figure 2) over time. The corresponding uptake flow rate (G_bl,up_ [kg s^-1^]) was also determined, which is the slope of the curve. Using this parameter, it was possible to calculate the flux across the skin into the blood (g_bl,up_ [kg m^-2^ s^-1^]).

The maximal fentanyl concentration in the stratum corneum (c_sc,max_(t) [kg m^-3^]) was also quantified. This quantity is important because concentrations of specific drugs that are too high may induce irritation [15]. With these quantities of stored and transported amounts of drugs, the mass balance of the drug can be written as:

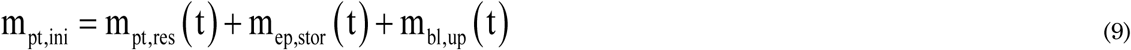

Out of the drug uptake profiles, *m*_*bl,up*_(t), the uptake kinetics were assessed by defining the fractional drug release of the patch (*Y*_*bl,up*_):

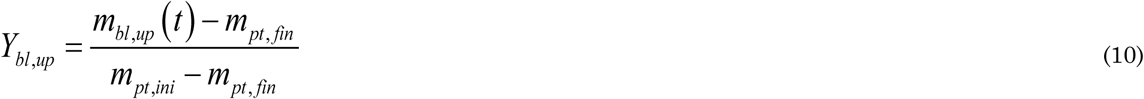

where subscripts *ini* and *fin* represent the initial and final drug amount in the patch when the patch is depleted, where *m*_*pt,fin*_ =0. m_*bl,up*_*(t)* represents the amount of the drug taken up by blood flow. From the definition of *Y*, the half-uptake-time (HUT, *t*_*1/2*_) can be calculated as the time required to take up half of the initial drug amount in the patch via the skin. HUT is a useful parameter to characterize and compare the release behavior of the patch, as a single value can be used to characterize the uptake kinetics.

## 3 RESULTS and DISCUSSION

### 3.1 Validation

The comparison with previous experimental data [21] in Figure 3 identifies how accurate our mechanistic model predictions on drug uptake through the epidermis are for two different initial drug concentrations in the reservoir. For both initial concentrations, the simulations are in good agreement with experimental data, i.e., within the error bars, during the first 27 h with a root mean square deviation (RMSD) of 0.13 and 0.14 µg cm^-2^ h^-1^ at 60 kg m^-3^ and 80 kg m^-3^, respectively. For the remaining two days, there are larger deviations. The experimental decrease in the flux exceeds that of the simulations. Model improvements that could enhance the accuracy are discussed in section 4.1. Note that the differences in these predicted fluxes and the cumulative drug amount that was taken up by blood flow between the 3D (cylindrical, Supplementary Material 2) and unidirectional (1D) transport models were < 0.4%. Therefore, a 1D transport model can be used as a viable alternative for the 3D model.

**Figure 3.**
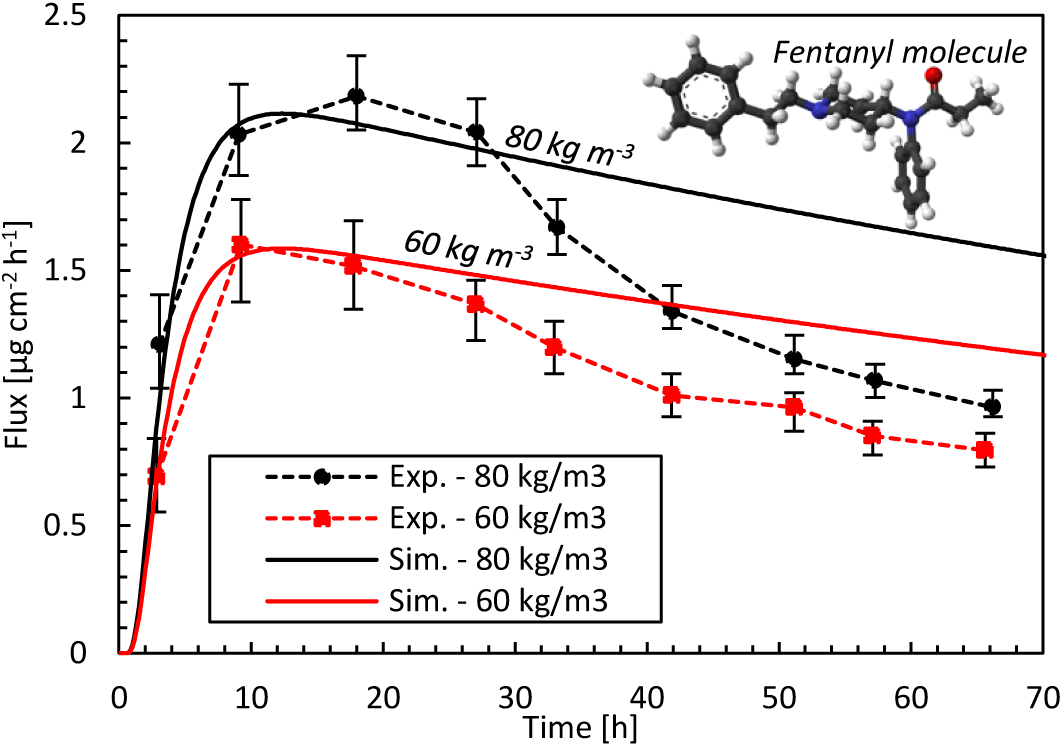
Drug fluxes of fentanyl at surface 1 (g_bl,up_), so leaving the epidermis, from experiments (Exp.) and simulations (Sim.) for two initial patch concentrations (60 kg m^-3^, 80 kg m^-3^) as a function of time. The fentanyl molecule is illustrated as well, where the oxygen is depicted in red, carbon in black, hydrogen in white, and nitrogen in blue.

### 3.2 Comparison with commercial TDDS

The drug uptake from the simulated transdermal patch (base case) is compared with that of commercial TDDS in Figure 4. From our simulations, we obtain a peak uptake rate by the blood of 23 µg h^-1^ after 7-10 h (Figure 4). This uptake decreases to 18 µg h^-1^ after 72 h for a 16 cm^2^ patch. The delivery rate is rather constant, but there is a slight decline due to the reduction of reservoir concentration, which is the driving force for drug transport. Commercial patches, however, only report the targeted steady-state value that does not reflect this slight decline in flow rate, because these TDDS are designed to deliver the drug at a nearly constant rate. Our results lie between the performance of Durogesic® DTrans® 12 µg h^-1^ and 25 µg h^-1^ patches, for example, which have a surface area of 5.25-10.5 cm^2^. In summary, our mechanistic model produces similar uptake rates and kinetics as commercial products. Combined with the validation study results, this tool is reliable for product design and optimization.

**Figure 4.**
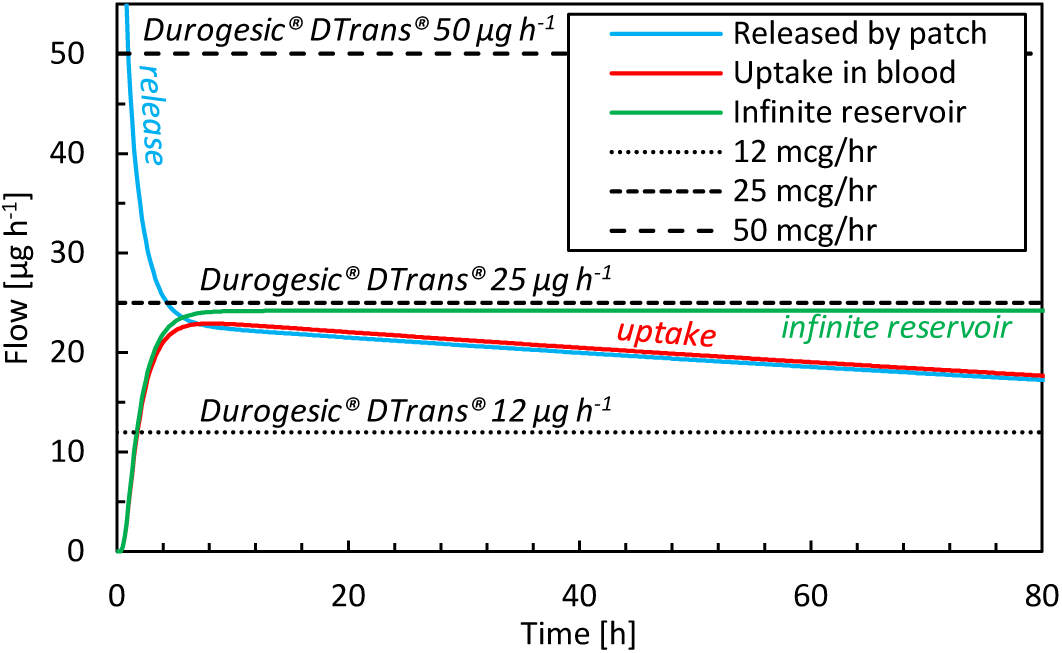
Drug flow as a function of time for drugs released by the patch (g_pt,rel_), for drugs taken up by the blood flow (g_bl,up_) for the base case and commercial fentanyl patches. The results of an infinite reservoir are also shown

### 3.3 Sensitivity analysis to model input parameters

The relative sensitivity of the flux across the skin into the blood (g_bl,up_) to the input parameters is shown in Figure 5a. These results quantify the parameters that affect the most the predicted drug uptake, and how this sensitivity changes over time. The total amount of drugs taken-up after a certain period is shown in Figure 5b. A sensitivity value *S*_*U,Xj*_ of 1 implies that the impact on the solution (U_Xj+ΔXj_ - U_Xj_) is more significant than the prescribed disturbance (1%, ΔX_j_) of the model input parameters (Eq.(7)).

**Figure 5.**
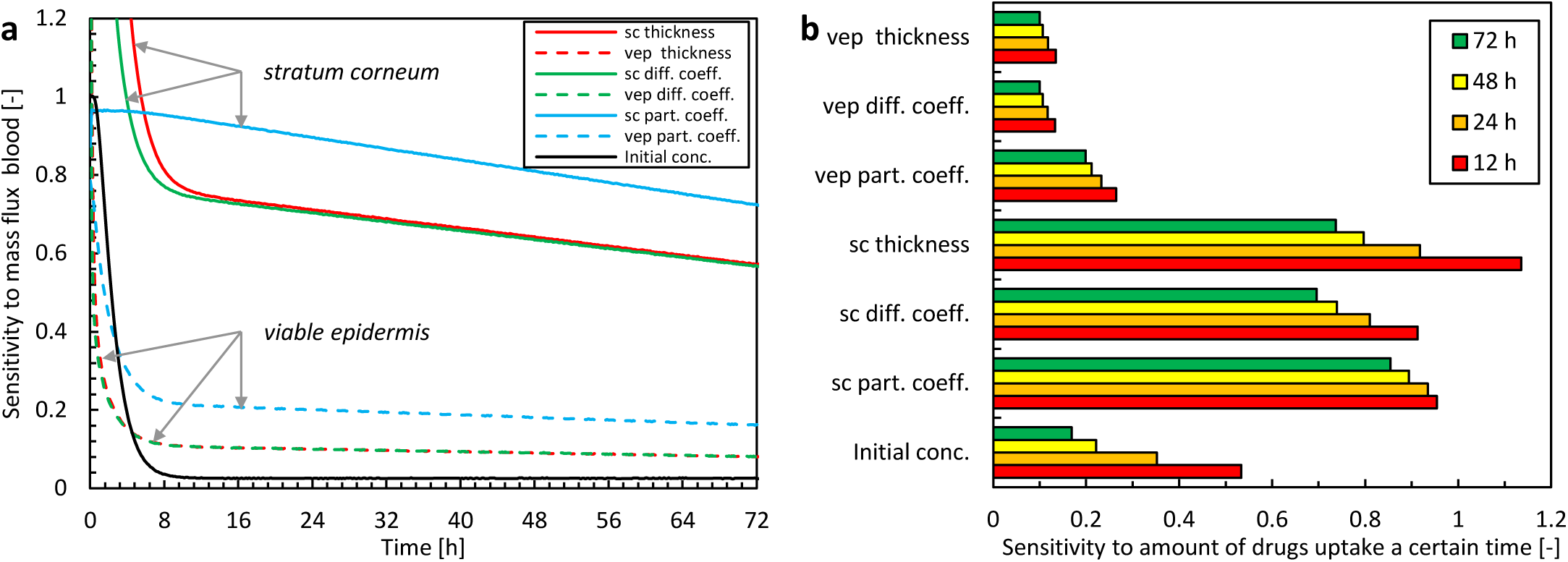
Relative sensitivity to different model parameter inputs for the mass flux taken up by the blood (U(t) = g_bl,up_) as a function of time (a) and the total amount of drugs taken up after 12-24-48-72 hours (U(t) = m_bl,up_) (sc: stratum corneum, vep: viable epidermis, diff. coeff.: diffusion coefficient, part. coeff.: partition coefficient, conc.: concentration).

Concerning the magnitude of the sensitivities, the data show the largest sensitivity for the model parameters that describe the stratum corneum. This finding is not surprising given that this epidermal sublayer has the most significant resistance to drug transport, and exhibits the skin’s primary barrier function. The partition coefficient has the largest overall impact of all model parameters. The thickness and the diffusion coefficient have similar sensitivities due to their comparable role in the conservation equation (Eq.(1)).

There is a distinct temporal sensitivity to the different model parameters, especially the thickness and diffusion coefficient of the stratum corneum and the initial drug concentration in the reservoir. This temporal sensitivity originates from the transient nature of the uptake process. Initially, loading of the drug in the skin occurs (first 10 h), which involves drug storage and transport through the skin. After this initial period, the sensitivity is more constant over time. The highest temporal sensitivity occurred for the initial drug concentration, with a sensitivity to the mass flux that reduces to only a few percent after the first 10 hours (Figure 5a). For designing drug delivery systems and therapy, the initial drug concentration in the patch is an important factor, and also comes into play when replacing the patch.

In summary, uncertainty in model parameters impacts the solution. Still, this effect is smaller than the disturbance itself (i.e., *S*_*U,Xj*_ <1), except during the very early stage of drug uptake (< 24 h). Nevertheless, the solution is particularly sensitive to the model parameters of the stratum corneum, especially the partition coefficient, so these parameters must be known as accurately as possible.

### 3.4 Drug release and uptake in the skin

#### 3.4.1 Release and uptake kinetics

The drug release and uptake kinetics from the patch into the skin are presented in Figure 6 for the base case, specifically by analyzing the contributions to diffusion and storage processes (including partitioning). These data display what the temporal delay is between drug release and uptake, and how much drugs are stored in the different skin layers over time. After drug release initiation, it takes roughly 30 min before the drug diffused through the 85 μm thick epidermis and can be started to be taken up by the blood via the capillaries in the dermis. In parallel to diffusive transport, drugs accumulate in the skin during the first 6-7 h. Despite its small thickness, the stratum corneum stores approximately 99% of the drug in the epidermis, because of its much larger capacity due to partitioning compared to the viable epidermis (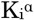, Table 3). Equilibrium occurs after this initial period because the skin’s capacity to store drugs is reached then. After that, the drug primarily diffuses through the skin, and the stored amount remains rather constant. The stored drug amount in the skin is, however, only a fraction of the total drug amount initially present in the patch, with a maximal value of 0.16 mg, or 2.4% of the initial content. This low value indicates that the drug bioavailability from the model was > 97.6%. Due to the decreasing drug concentration in the patch (a finite reservoir), the uptake flux slowly decreases over time, simply because the concentration gradient decreases.

**Figure 6.**
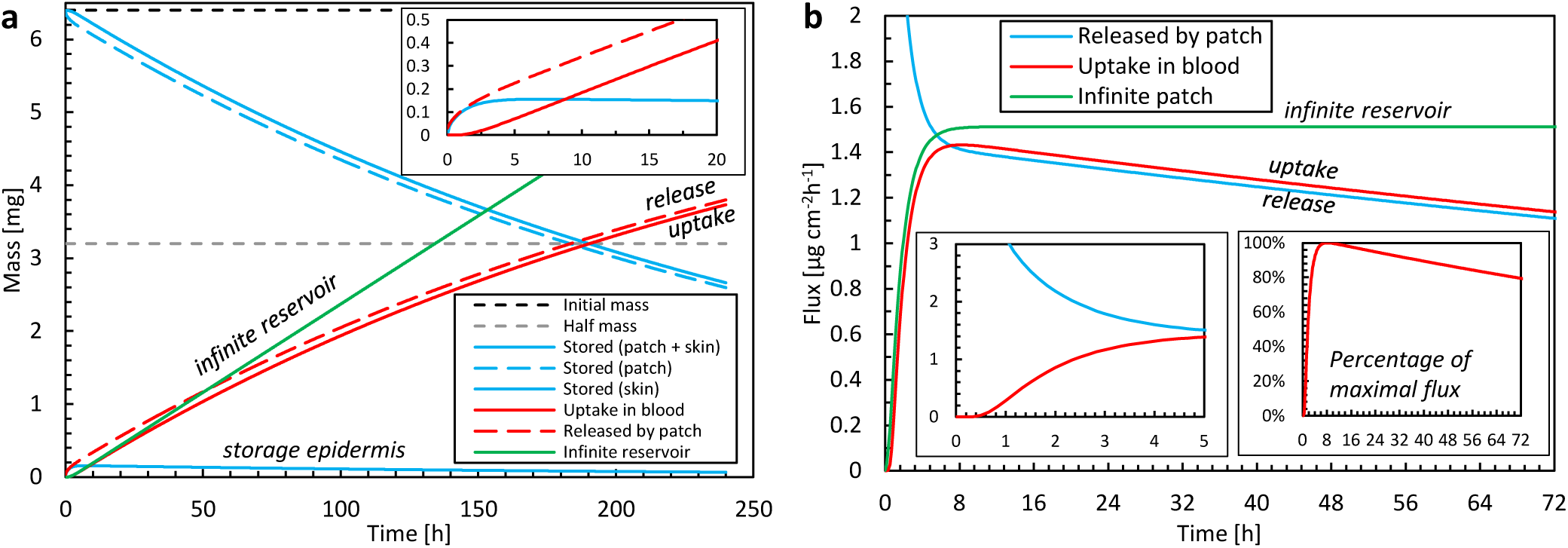
(a) Amount of drug stored in the epidermis (m_ep,stor_(t)) and patch (m_pt,res_(t)), and cumulative amounts of drugs released by the patch m_pt,rel_(t) and taken up by the blood flow m_bl,up_(t) as a function of time for the base case. The initial amount of drugs is also shown. (b) Corresponding flux released by the patch into the epidermis (g_pt,rel_) and taken up by blood (g_bl,up_) as a function of time, where one insert shows the percentage for the maximal value. The results of an infinite reservoir are also shown.

The patch is depleted by half of its initial amount after 190 h (almost 8 days). Typically, fentanyl patches are replaced every 72 h or 3 days (e.g., Table 2). After this 72-h application period, 77% of the initial drug amount is still present in the patch for the base case. However, transdermal patches are designed to deliver the drug at a controlled rate to achieve a constant blood plasma concentration rather than delivering the entire amount of drug and depleting the reservoir [1]. Therefore, the concentration in the patch cannot reduce too much, as this would imply the driving force for drug transport, i.e. the concentration difference, would become too low. As such, a significant drug concentration should still be present when removing the patch, to guarantee a rather constant drug flow [60]. Also, in our validation study, only 31% of the initial drug amount diffused through the skin after 72 h. Larsen [6] reported absorbed amounts below 20%.

For comparison, the results of an infinite reservoir with high diffusive drug transport are also shown in Figure 6. As expected, a constant flux is reached after an initial uptake period. In other words, there is a linear increase in the drug amount taken up with time. Ideally, TDDS are designed to deliver drugs at a constant rate. The resulting constant fentanyl flux (1.5 μg cm^-2^ h^-1^) is similar to the range reported for commercial patches (Table 2; [2]), namely 1.7-3.0 μg cm^-2^ h^-1^. The finite reservoir deviates from this constant rate. After 72 h, the flux is only 80% of the maximum (insert in Figure 6b). The infinite reservoir overpredicts the amount of drug uptake by 17% after 72 h compared to the finite reservoir (base case), and this difference is significant.

Additionally, Supplementary Material 3 shows that the differences with a 3D model are limited, and the anisotropic transport properties between transverse and longitudinal direction have a limited impact on the evaluated patch width.

#### 3.4.2 Spatial and temporal resolution

We compare the high spatiotemporal resolution data from the simulations against typical data obtained in TDDS experiments. This endeavor identifies additional insights and benefits gained from mechanistic modeling. In experiments, drug uptake is typically measured using a Franz diffusion cell at discrete time intervals (e.g., every 5 h) and subsequent measurements of the samples via HPLC [6], [21]. From the measured drug amount taken up (μg) during the sampling interval (h) and the patch’s surface area (cm^2^), the time-averaged flux through the entire skin sample can be calculated (μg cm^-2^ h^-1^). Even if the total amount of drug taken up is correct, such temporal averaging masks local flux peaks in time and the associated maxima in concentrations. This information is essential because concentrations of specific drugs that are too high may induce skin irritation [70], [71]. The discrepancies induced by such temporal averaging are quantified in Figure 8, as derived from our simulation data. The average flux for different time-averaging intervals is calculated by grouping our simulation results over specific intervals because these can be derived from the fluxes at each point in time. Averaging intervals > 5 h substantially underestimate fluxes by approximately 20%. However, this accuracy strongly depends on the process kinetics during that timeframe. Simulations also allow researchers to mimic experimental conditions in a deterministic way, without suffering from biological variability and the resulting uncertainty.

**Figure 7.**
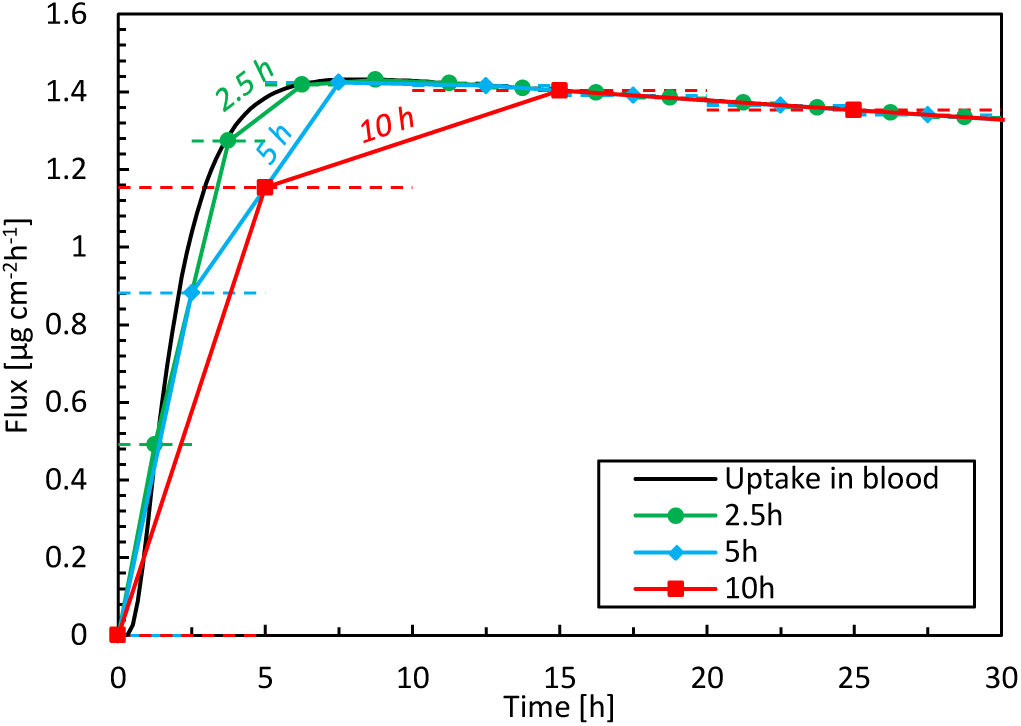
Flux taken up by the blood as a function of time for three time-averaging intervals, based on simulation data. The horizontal dashed lines indicate the time range over which the averaging took place, i.e., 2.5 h, 5 h or 10 h.

**Figure 8.**
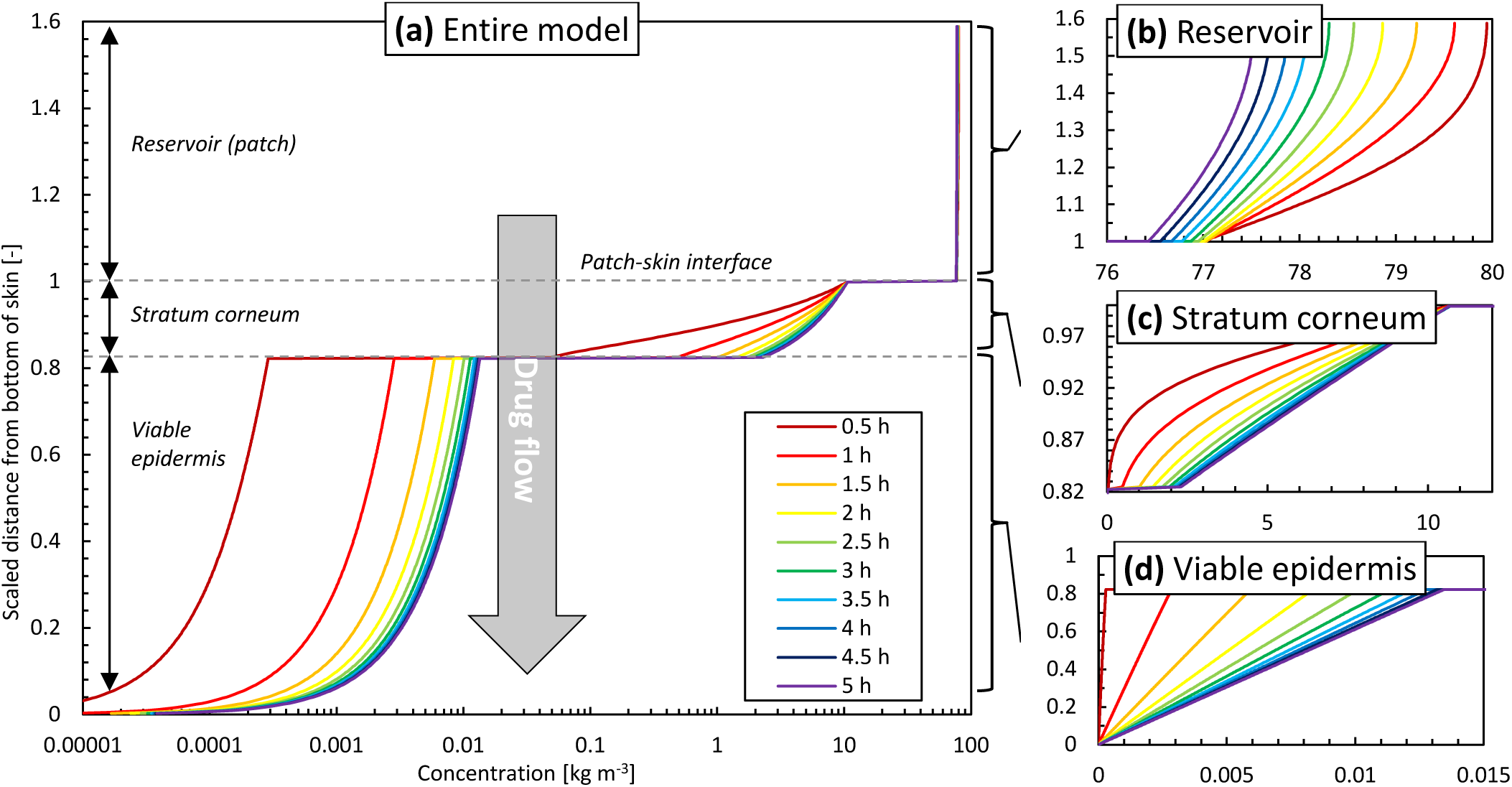
Concentration profiles at different points in time as a function of the height of the model (z-direction, scaled with the epidermal thickness) in the vertical centerline (z-axis) of the patch for the base case: (a) entire model (logarithmic scale), (b) reservoir, (c) stratum corneum, (d) viable epidermis.

To further illustrate the added spatiotemporal insights on drug transport obtained by simulations, the vertical concentration profiles through the patch and skin are shown in Figure 8. The experimental counterpart to obtain such profiles would be tape stripping [72]. This process, however, only extracts the profiles at a single point in time, where typically steady-state conditions are targeted. The obtained 1D spatial resolution by tape stripping is in the micron range, but is strongly dependent on the anatomical site, age, stratum corneum thickness, number of cell layers, and corneocytes, among others [72]. Contour plots of concentration and potential from the simulations are shown in Figure 9. Partitioning is indicated by the much higher concentrations of the moderately lipophilic drug in the stratum corneum compared to the viable epidermis. The discontinuities at the interface of the skin-patch layer are visible and contrast the continuous distribution of the dependent variable drug potential *ψ*^*α*^ (Eq. (4)) through all layers. These large concentration jumps challenge the numerical stability of the simulation, which is why the conservation equations were solved against the drug potential instead of drug concentration in this study.

**Figure 9.**
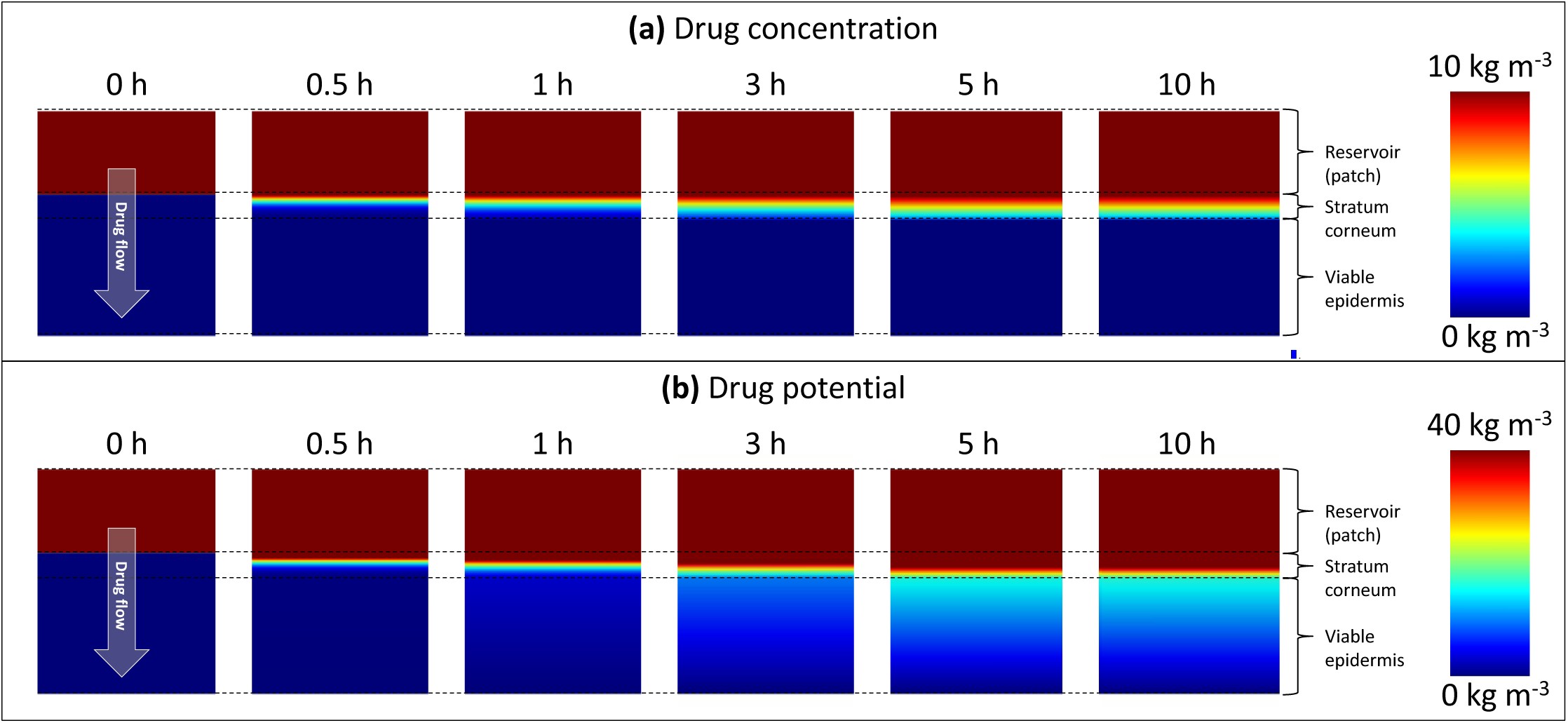
Color contours of (a) drug concentration (c_i_^α^) and (b) drug potential (ψ^α^) in the drug reservoir and epidermis for different points in time. Different scaling is used in (a) and (b) to improve clarity. Note that the maximal range (80 kg m^-3^) is not depicted.

After initially loading the epidermis with the drug, a quasi-steady-state diffusion process develops, with the typical linear concentration distribution over each epidermal layer (Figure 8c and d). The concentration reduction in the patch (Figure 8b), however, still induces a slight temporal shift in the concentration profiles in the epidermis (Figure 8c, d).

#### 3.4.3 Removing patch after 72 hours

The simulations enable us to analyze what occurs within the epidermis when the fentanyl patch is removed after the recommended 72-h period. This analysis is possible because simulations can quantify volume-averaged drug contents as well as surface-averaged fluxes. In Figure 10, the drug uptake amount in the blood, the corresponding flux, and the drug storage in the skin are shown as a function of time. A very small drug amount was stored in the skin (0.12 mg after 72 h). Because only this amount can further diffuse into the blood once the patch is removed, the drug amount that is taken up steeply decreases after removal. However, it still took approximately 24 h before this residual amount diffused from the skin to the blood. This effect is more pronounced for drugs that can be stored in higher amounts in the skin, namely those with a larger drug capacity 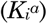.

**Figure 10.**
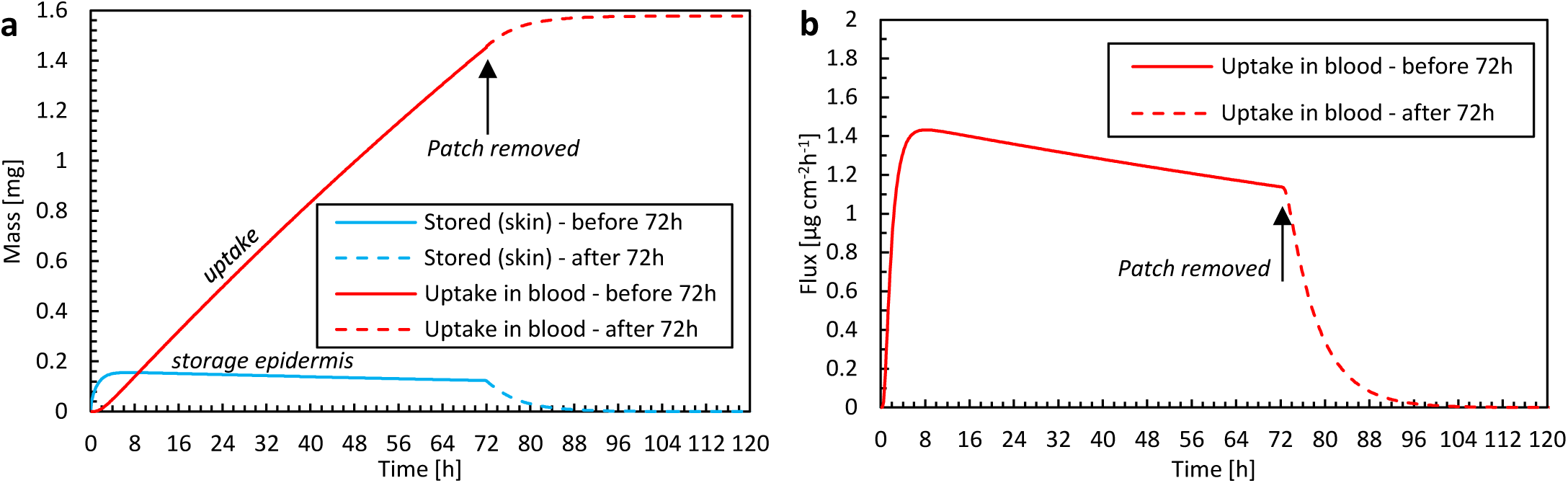
(a) Amount of drug stored in the epidermis (m_ep,stor_(t)) and the cumulative amount of drugs taken up by the blood flow m_bl,up_(t) as a function of time for the case where the patch is removed after 72 hours. (b) The corresponding flux that is taken up by blood (g_bl,up_) as a function of time.

### 3.5 Patient-induced variability for anatomical location

The impact of the anatomical location of the drug reservoir on the human body was also investigated. The absorption of drugs through the skin differs in different body locations due to a different thickness of the stratum corneum or the presence of the hair follicles [73]. In Figure 11, the amount of drug uptake is given for three-body sites that differ in terms of epidermal thickness. The HUT (section 2.5) is also indicated for each body location, as calculated based on the curves with average thickness. The HUT for the base case was 190 h.

**Figure 11.**
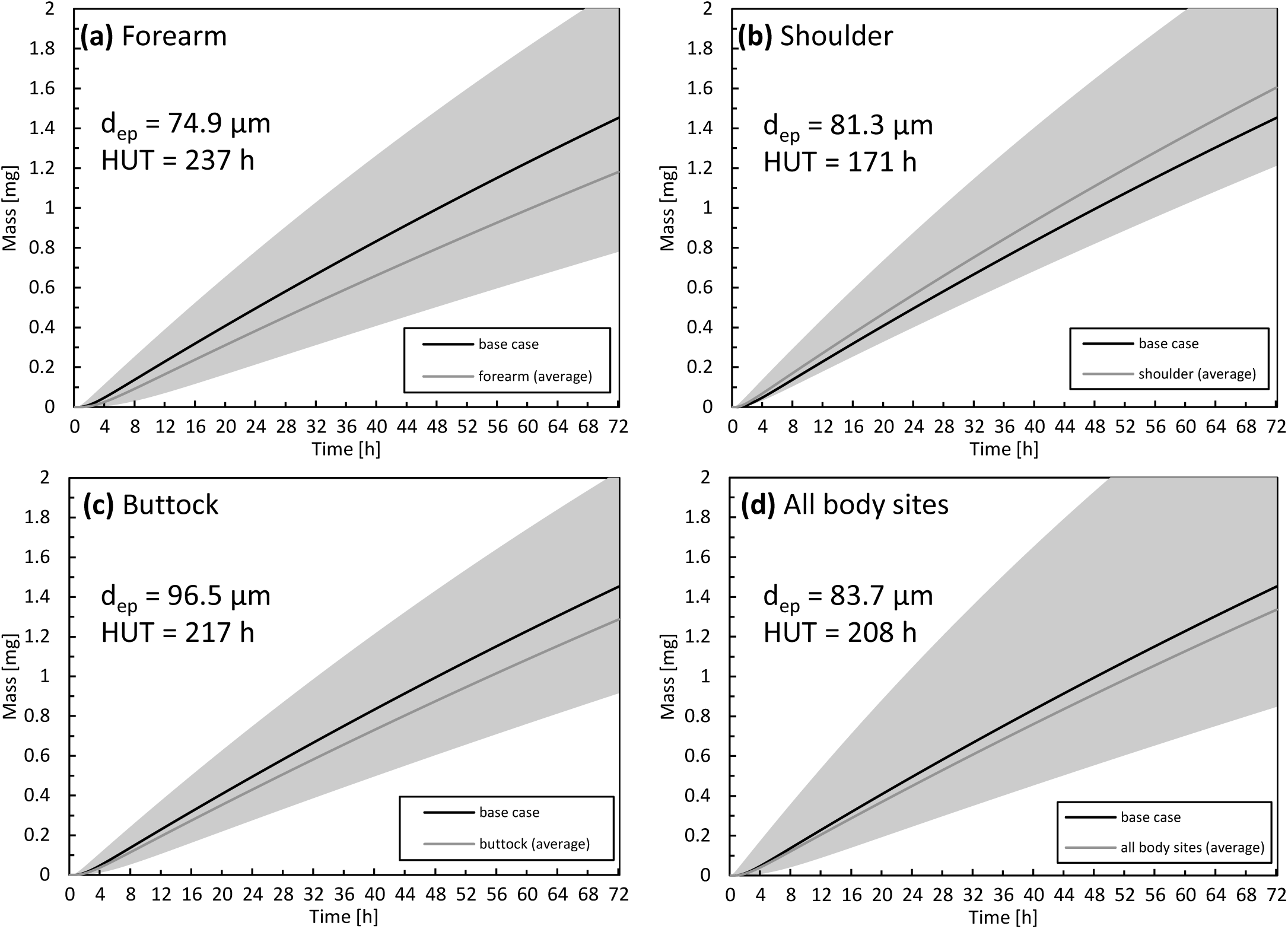
Cumulative amount of drug taken up in the blood as a function of time for the forearm (a), shoulder (b), buttock (c) epidermal thickness averaged for all body sites (d). For comparison, the results of the base case are also depicted. The range corresponding to the simulated average thickness values +/- 2 x standard deviation (µ_i_ -/+ 2σ_i_) is depicted in gray to quantify interpatient variability via the 2.5% and 97.5% quantiles. The thickness of the epidermis (d_ep_) and half-uptake time (HUT) are indicated as well.

The largest drug uptake was observed on the shoulder, while the smallest was obtained for the forearm. The difference in drug uptake (mg) between these two body sites was 36% (relative to that of the forearm) after 72 h. The results for the average epidermal thickness over all the body sites agree with the base case. Interpatient variability in the stratum corneum and epidermis thickness directly manifested itself in drug uptake rates. For the forearm, the variation in total drug uptake (mg) over the simulated range (µ_i_ -/+ 2σ_i_) is 113%, and this variation is not evenly distributed across the average value. Clinically, this finding implies that specific patients can have blood serum concentrations that can be too high (toxic) or too low to be effective for that specific patient. This interpatient variability for a specific anatomical location induces significant variability on the drug uptake results. This spread makes it challenging to distinguish significant differences between anatomical locations in clinical experiments. In contrast to the present study, specific previous studies reported limited differences in fentanyl uptake between the anatomical location [6], [7].

### 3.6 Patient-induced variability for age

The impact of patient age on drug uptake kinetics was investigated by calculating variations in the stratum corneum thickness with age via Eq. (8). In Figure 12, the drug uptake and flow rate are given for different age groups (as previously defined [10]). There is a 26% difference in the total drug amount taken up after 72 h between an 18- and a 70-year-old patient (relative to the 18-year old patient). This finding suggests that if a patient applies the same fentanyl patch now and 52 years later, this patient will receive 26% less drug when s/he is older; therefore, the patch will be less effective. When examining flow rates, one could even consider performing therapy using a patch with a higher dose and/or a larger surface area. The current mechanistic model can even be used to define an age-specific dosage systematically.

**Figure 12.**
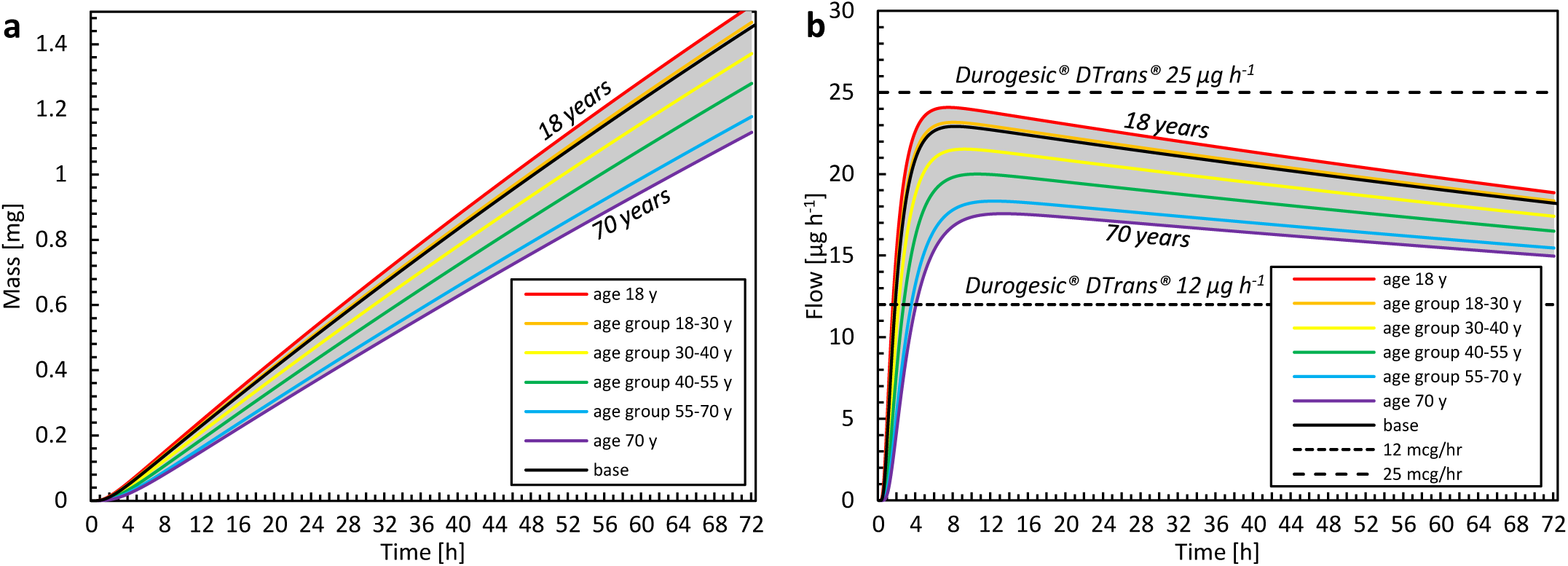
(a) Cumulative amount of drug taken up in the blood as a function of time for different age categories. The results of the average age of each group are shown, as well as the minimal and maximal ages. (b) Drug flow taken up by the blood as a function of time for different age groups and commercial fentanyl patches. For both a and b the gray band indicates the entire range of ages that is evaluated (18-70 years).

Furthermore, the time before the minimal effective concentration is reached in the blood differs with age. As an example, it took 23 h to uptake 0.5 mg for an 18-year-old patient, whereas it took 9 h longer when the patient was 70 (Figure 12a). Our simulations showed that, with aging, the patch delivered drugs more slowly and less potently. Mechanistic simulations enable researchers to quantify this difference deterministically and theoretically, without introducing additional statistical uncertainty concerning interpatient variability.

### 3.7 Impact of contact surface area of the reservoir

We explored how the reservoir width, and thus the contact surface area affected the released flux. For normal TDDS, the reservoir is much wider than the epidermal thickness, which is the longitudinal transport pathway for the drugs. This phenomenon leads to unidirectional transport. Hence, the 3D edge effect induced by transverse diffusion at the edges of the patch is negligible. This examination aimed to identify whether a reservoir with a smaller contact surface area released drugs faster and at a higher rate than a standard patch. Since transport will occur in 3D in the case of patches with a smaller contact area, this transverse diffusion could induce higher fluxes. In Figure 13, the released flux (surface 2) is shown as a function of time for all reservoir surface areas for finite and infinite reservoirs. In Figure 14, the released flux at equilibrium (for an infinite reservoir) is shown as a function of reservoir width (L_pt_), where the reservoir width is made dimensionless with the epidermal thickness (*d*_*ep*_ = 85 μm).

**Figure 13.**
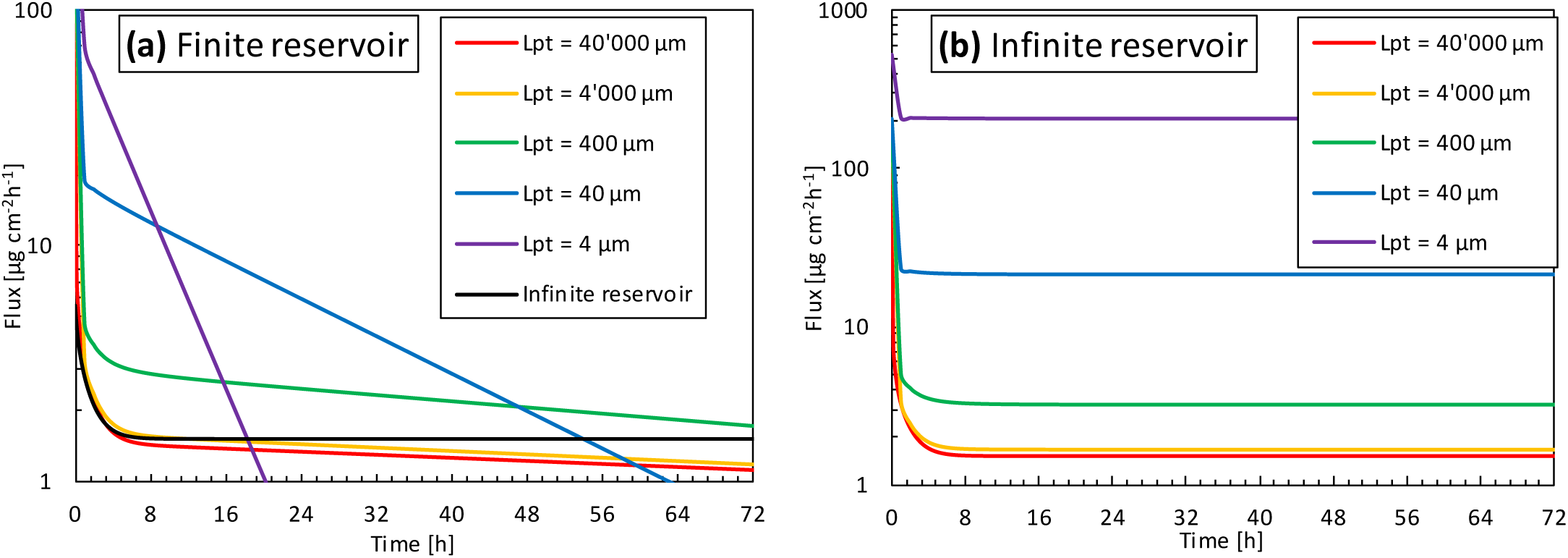
Flux released by the patch into the skin (g_pt,rel_, surface 2) as a function of time for different reservoir sizes (L_pt_), so contact surface areas, for (a) finite reservoir, (b) infinite reservoir.

**Figure 14.**
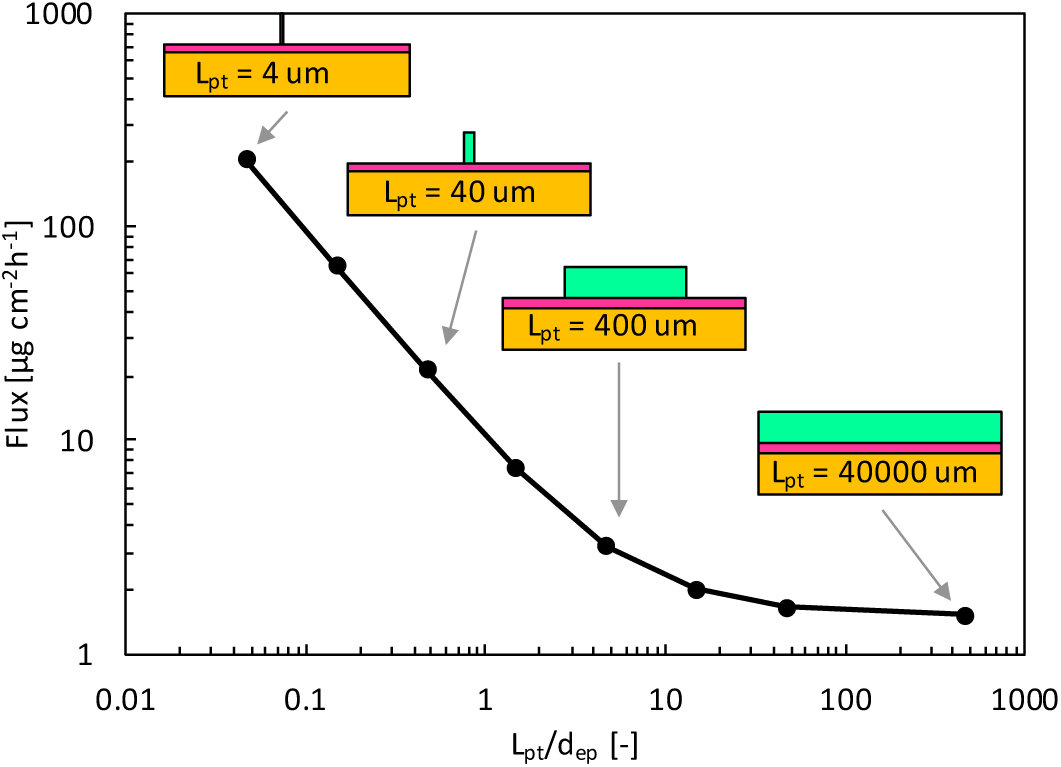
Flux released by the patch into epidermis after 72 hours (g_pt,rel_) for the infinite reservoir as a function of the reservoir size (L_pt_), so contact surface area, scaled by the epidermal thickness (d_ep_). In the schematics, dimensions of the patch and skin thickness are to scale, but the skin width is not to scale.

The flux leaving the reservoir is dependent on the size so contact surface area of the drug reservoir in contact with the skin, for both finite and infinite reservoirs. For the finite reservoirs, the depletion of the smaller reservoirs causes the flux to decrease sharply over time. This depletion can, however, be mitigated by increasing the thickness of the reservoir (*d*_*pt*_) or by connecting all small reservoirs to a large bulk reservoir. The infinite reservoirs evolve to a steady-state, a condition that is more convenient for comparison of the reservoir contact surface area. Once the size (L_pt_) enters the submillimeter range, or the patch size goes below approximately 10 x d_ep_, the flux increases to more than double of that of a conventional patch (base case). This phenomenon occurs because the drug can diffuse in three dimensions instead of predominantly one direction for the large reservoirs.

Furthermore, the transverse diffusion coefficient of the stratum corneum is a few orders of magnitude larger than the longitudinal one ([26]; Table 3). Thereby, for smaller contact surface areas, longitudinal and transverse diffusion occur, a process that induces a higher flux at the contact interface. This edge effect is illustrated in the contour plots presented in Figure 15. For a reservoir size (L_pt_) of 40 μm, the release rate increases by a factor of 20 at equilibrium for an infinite reservoir compared to the base case. For L_pt_ = 4 μm, the increase was 200-fold. Note that these factors decrease to 3 and 18, respectively, when the transverse diffusion coefficient would just equal the longitudinal one (results not shown). This data implies that transverse diffusion is a key parameter for the large observed differences, partially due to the higher transverse diffusion coefficient of the stratum corneum layer, especially when the reservoir is not very wide compared to the skin thickness. Note that for small patches, drug storage in the skin will delay uptake into the blood, because it takes longer to saturate the skin sublayers due to their lower total flow rate (µg h^-1^). It would take longer to reach an uptake equilibrium (g_bl,up_), namely approximately 35 h for L_pt_ = 4 μm versus 15 h for L_pt_ = 40,000 μm. The amount stored in the skin is, in all cases equal to, or smaller than the base case. Note that the smallest reservoirs are of the same size as the corneocytes (Figure 1) but still much larger than the lipid bilayer thickness (approximately 10^1^ nm [74]). As such, the drug concentration contours could depend to some extent on where the reservoir is precisely placed, relative to the corneocyte or lipid bilayer at the surface. This can be visualized with a mesoscale model (Figure 1). However, in the current lumped approach with anisotropic diffusion in the SC layer, we receive an average drug uptake profile. This is justified if we assume that for a complete patch, the reservoirs are randomly located on the skin by which the average uptake we simulate is still representative. Note that the trend we see for these very small reservoirs is already present for the reservoirs that are much larger than the corneocytes (Figure 14). This implies that smaller reservoirs progressively take more benefit more of the transverse diffusion, compared to larger reservoirs, to increase the drug uptake flux.

**Figure 15.**
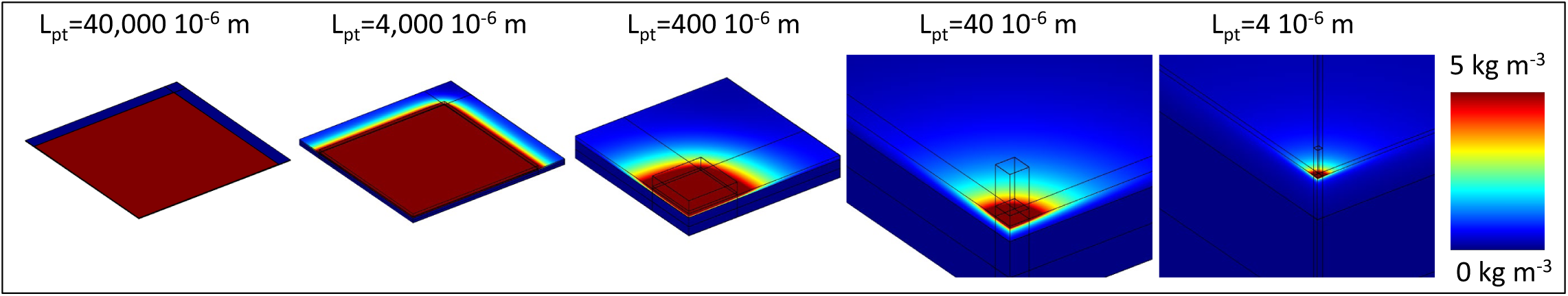
Color contours of drug concentration over the skin for different sizes of the reservoir for simulations with an infinite reservoir after 72 hours, so when a steady-state is reached. Note that the maximal range is not depicted (80 kg m^-3^), and only one-fourth of the patch-skin system shown due to symmetry.

## 4 OUTLOOK

### 4.1 Mechanistic modeling

The current state of the art in mechanistic modeling of TTDS was summarized in Table 1. Such mechanistic modeling provides several advantages compared to the analytical solution of the diffusion-driven drug uptake process [21], [75]. Such analytical solutions enable to calculate drug dose taken up for several drugs, based on their diffusive properties of the skin. Mechanistic modeling is, however, required to target more complex situations, for example, when considering the skin as a multi-layer structure (stratum corneum, viable epidermis) or when the patch is replaced so if the boundary conditions change over time. Compared to the current study and state of the art (Table 1), further advancements should be pursued to enhance the realism and accuracy of transdermal drug delivery predictions in terms of (1) the modeled transport processes, (2) the targeted drug delivery system, (3) numerical modeling, and (4) the model parameters. These future targets for model development are detailed below.

#### 4.1.1 Modeled transport processes

Concerning transport processes, the physical adsorption of molecules or chemical binding should be included to enhance accuracy. For fentanyl, bioavailability through the skin is very high (e.g., 92% in [60]) but not equal to 100%. Thus, some of the fentanyl does not reach the systemic circulation, due to absorption in the epidermis or by chemical changes. Including these mechanisms in mechanistic models is not yet a standard practice [76], [77]. The current mechanistic model for transdermal drug delivery is built up for first-generation systems [78]. Thereby, in addition to fentanyl, it can also be calibrated for lipophilic drugs with a small molecular weight, for example, ibuprofen[79], buprenorphine, rotigotine, and rivastigmine [80]. For next-generation systems [55], [81] that enable the delivery of larger molecules like insulin, additional processes would need to be included in the current mechanistic model. These systems aim to enhance skin permeability by increasing the driving force for drug uptake via chemical permeation enhancers, iontophoresis, or by disrupting the stratum corneum using microneedles or thermal ablation [55].

Furthermore, when including the dermis in the model [19], [35], [36], which is implicitly done when modeling skin thicknesses in the millimeter range, blood flow (in addition to diffusion) must be modeled. Modeling the dermis without drug extraction by blood flow will underpredict the drug uptake rate. The impact of including the dermis in the modeled system configuration is illustrated in Supplementary Material 4 (Figure S2), where the impact of different dermis thicknesses on diffusive drug transport is illustrated, so without modeling blood flow. Due to the relatively large dermis thickness and volume, modeling only diffusive transport overpredicts the amount of stored drug and transverse spreading found in specific studies [36]. The impact of tight junctions [47] on diffusion was not explicitly modeled but should be considered in the future.

Additionally, skin swelling or shrinkage, for example, driven by changes in skin hydration, could be included [82]. For an infinite reservoir under steady-state conditions, where Fickian diffusion would predict a constant flux, swelling will introduce a time-dependency into the flux [1]. The swelling or shrinkage of the patch could also be considered [21]. Finally, diffusion and partition coefficients that are a function of drug concentration (rather than constant values) should be used, especially if there are large variations for the drug of interest. For example, when increasing the Azone concentration from 0 to 9 wt%, the partition coefficient changed from −4.0 to 2.1 [25]. This alteration significantly changes its equilibrium distribution. Including all of the aforementioned physical processes in the model is not always required as they don’t all play critical roles for each drug molecule. Their importance should be assessed on a case by case basis.

#### 4.1.2 Drug delivery system

Concerning the modeled delivery system, finite reservoirs should always be preferred over infinite reservoirs, which are currently still commonplace (Table 1). For finite reservoirs, the gradient, therefore the driving force, will decrease, a phenomenon that makes delivery at a constant rate more challenging (Figure 6). For this reason, controlled drug delivery systems were designed to alleviate the decreasing concentration gradient (e.g., multilayer Deponit® system [1], [44]). Future models should also include the dermis. This inclusion is essential to evaluate the drug share taken up by the blood versus the amount of the drug that diffuses into and is stored in the thick dermis. This factor will affect total bioavailability and uptake kinetics. In this study, the storage in the dermis was assumed small compare to the amount of drug taken up via the dermis in the blood flow, due to the large bioavailability of fentanyl. If the dermis is included in the computational model, it is essential to include the blood flow in capillaries and vessels and to adjust this blood flow as a function of patient age and activity level, among others [83]. Another reason to include the dermis is that the current boundary condition imposed at the viable epidermis (surface 1), namely a zero concentration, has a certain limitation. This condition is valid in experimental setups with Franz diffusion cells, but it can be disputed for a real human tissue. Here, the concentration profile at the viable epidermis will result from the trade-off between diffusion and uptake by the blood flow. Finally, the mechanistic model should be linked to a pharmacokinetic model that relates the uptake amount to the metabolization process in the body to obtain the final blood plasma concentration.

#### 4.1.3 Numerical modeling

Concerning numerical modeling, there were large discontinuities in concentration over the skin layers due to partitioning. To improve numerical stability and accuracy, it is advised to solve for a dependent variable other than concentration, as was performed in the present study using drug potential.

#### 4.1.4 Model parameters

Concerning the model parameters, the diffusion and partition coefficients are rarely measured explicitly before modeling using a separate *in vitro* experiment. Instead, data from the literature are utilized, often even from other drugs with similar molecular weight and lipophilicity [26]. Alternatively, data are also fitted [21] or corrected [33] to match experimental data. Obtaining a good agreement, in this case, is not surprising and can mask missing physical processes within the model. Therefore, one cannot claim the model is validated, but rather it should be considered calibrated. This procedure to obtain model parameters is not necessarily discouraged, but one cannot use the same dataset for model fitting and experimental comparison. Selzer et al. [36] recently fit model parameters to *in vitro* experiments and compared the calibrated model to a set of *in vivo* experiments. This approach is certainly viable, but the resulting model parameters still led to differences in the experiments.

### 4.2 Patient variability and personalization

The impact of aging on transdermal drug delivery was considered by changing the stratum corneum thickness. For a 70-year-old patient, the drug taken up by the blood was lower, and the maximum peak flow of the drug occurred later in time than for an 18-year-old patient. This *in-silico* data analysis quantified the dose delivered for differently aged patients for a specific drug concentration in the patch. Moreover, the slight time shift in peak drug flow is also helpful for defining the time window where patients of different ages may be more susceptible to developing side effects. This information enables more precise monitoring and prevention of serious side effects. These data could be used to tailor devices for specific age groups or to develop devices that can monitor the drug flux and tailor it depending on the person’s age. This would be a step forward compared to current conventional transdermal fentanyl therapy. Where the initial dose (so patch size) is being estimated based on previous daily doses of oral morphine for the patient [84], with applying the patch transdermally and replacing it every 72 hours [40]. These features would allow a tailored treatment and a constant delivery rate below defined thresholds [81]. Additional age-related changes in the stratum corneum structure, such as a decrease with lipid content and its composition [85], lipid peroxidation [86], [87], or reduced hydration level [10], [88], were not accounted for in this study. There are several challenges with designing such tailored devices or therapy for transdermal drug delivery and implementing then into the clinics. The most straightforward solution that could be implemented the most swiftly would be to use conventional transdermal therapy based on existing fentanyl patches. The most optimal therapy with respect to the initial concentration in the patch, the amount of time it should be applied (currently 72 hours), and the location where it should be applied could be determined per age category. This means that a clinician could use the mechanistic model to decide which patch to use (e.g., Durogesic® DTrans® 12 µg h^-1^ or 25 µg h^-1^), how long to use it (72 hours or shorter/longer) and where to place it on the body.

### 4.3 Reservoir design and contact area

From our results, multiple smaller isolated reservoirs with the same total exposed surface area will likely release a drug at a higher rate than a single reservoir. This finding suggests that drug uptake could be enhanced only by patch design, namely by taking advantage of the transverse drug transport in the epidermis and particularly in the stratum corneum, without penetrating the stratum corneum layer with microneedles or ablating it. This finding could help in designing and individualizing TDDS by customizing the reservoir size and, thus, contact surface area to the patient. Identifying and quantifying this effect is only possible with simulations because accurately measuring these local fluxes over small reservoirs (e.g., 2 µm) would be very challenging experimentally.

Although the flux increases with smaller contact surfaces, the total amount of drug delivered (mg) will be lower, and these smaller reservoirs will also deplete faster (see Figure 13a). The latter is manifested by a rapidly changing magnitude of the flux over time for finite reservoirs. As such, for smaller reservoirs, it will be more challenging to deliver drugs at a constant rate, which is a key target for TDDS. However, the commercial application of such small reservoirs implies that multiple reservoirs could be integrated into a single patch with a larger thickness or a buffer reservoir to mitigate fast depletion. This design may mitigate the faster depletion and decreased rate, but this concept should be evaluated in more detail in the future to render these findings more conclusive.

Note that in this study, complete contact between skin and drug reservoir was assumed. This phenomenon implies that the material in the drug reservoir (e.g., a gel) is sufficiently deformable to ensure such perfect contact upon application. If the contact is not perfect, the skin roughness and the hydration level will also play a role in drug diffusion and delivery and thus must be explicitly considered [82].

### 4.4 Future use of mechanistic modeling in TDDS

A decisive factor in the future use of mechanistic modeling for TDDS design is to increase their efficiency. A key bottleneck in the workflow is typically the low availability of accurate and suitable model parameters and combining *in vivo* data from different studies, a factor that lowers the accuracy of the model solution [36],[89]. Currently, it is possible to obtain a satisfactory agreement between mechanistic model results and experimental data, but differences remain during the uptake process [19], [21], [27]. A better absolute agreement could be pursued by obtaining more accurate model parameters from detailed experimental measurements, for example, the *a priori in-vitro* parameter determination that was applied in one study [36], but even then some discrepancies with between simulations and experiments remained. However, if this time-intensive step needs to be performed for every new set up or drug of interest, *in silico* TDDS design will not be used beyond academic studies. This endeavor would require more time to obtain the model parameters for simulations rather than to perform stand-alone experiments. In this case, simulations would likely be of primary interest when a single drug is considered, and a very large parametric space is explored. An alternative for obtaining the model parameters from experiments would be inverse modeling.

Instead of pursuing a high precision in the predicted uptake kinetics [36], computer-aided engineering (CAE) in TDDS could, however, easily be used to probe for relative differences among systems, devices, drugs, or patients. For this purpose, model parameter data can be simply based on the literature, a feature that would make the entire modeling suite much swifter and more attractive. With this perspective, CAE has been used to push innovation in many other engineering fields, from modeling blood flow in vessels to the integrated design and construction of civil structures [90].

Finally, mechanistic modeling could become an essential component of fourth-generation controlled, feedback-induced TDDS. These TDDS enable multiple drug release rates and use feedback control based on measured biomarkers via wearable sensors to regulate drug delivery [81]. Currently, such wearable technologies are being developed to individualize treatment [81]. Wearable sensors measure biomarkers and thus, monitor the patient’s physiological condition, which is used to trigger the release of medication. Consequently, actuated transdermal drug delivery patches steer the drug release to provide the correct dose. The sensors and actuators are integrated in a closed feedback loop for better individualizing therapy to deliver the optimal rate at the right time to the correct body location. These TDDS rely on the measured patient’s response, i.e. the change of certain biomarkers, for control. There can be, however, a significant time lag between the drug release and the drugs reaching the systemic circulation. Mechanistic simulations could provide key complementary quantitative insight on the drug-release and percutaneous absorption kinetics for each patient, and help to better control fourth-generation TDDS.

In this context, a next step in these fourth-generation TDDS could be the use of digital twins. A digital twin is a virtual representation of the TDDS, which is linked to the real-world patient by sensor data of certain biomarkers. Digital twins can be used here for predictive modeling of the drug release and uptake in the human body, and can quantify the time lag, and account for it during control. Here, mechanistic modeling is an essential building block for digital twins of human organs.

## 5 CONCLUSIONS

Validated mechanistic modeling was used to gain new insights into transdermal fentanyl uptake. First, we quantified the changes in transdermal fentanyl uptake with the patient’s age and the anatomical location where the patch was placed. We also evaluated how much the drug flux can be enhanced by miniaturizing the drug reservoir size. Additionally, we obtained quantitative insights into the release and uptake kinetics of fentanyl transdermal drug delivery by analyzing drug diffusion, storage, and partitioning. The main findings are summarized below.

– Differences in drug uptake amount between anatomical locations of the drug reservoir on the human body of 36% after 72 h were found, but there was also strong interpatient variability.
– With aging, the transdermal drug delivery patch worked slower and less potently. An 18-year-old patient received 26% more drugs over the 72 h application period than a 70-year-old patient.
– Our proposed novel concept of using micron-sized drug reservoirs induced a much higher flux (µg cm^-2^ h^-1^) than conventional patches. Due to enhancing transverse diffusion in the stratum corneum layer, simply by changing patch design, a 300-fold increase in the drug flux was possible for a micron-sized patch. The identification of fluxes at the micron scale in a straightforward and accurate way was only possible *in silico*. With this concept, the reservoir surface area can be tuned and individualized for a specific patient or patient with a certain age category.
– For commercial patches, where the size is much larger than the epidermal thickness, drug transport was mainly unidirectional, and 1D models can be used reliably, assuming perfect contact between skin and reservoir.
– We showed *in silico* that sampling intervals in experiments > 5 h significantly underestimated peak drug fluxes, for example, by 20% when considering a 10-h interval.
– The role of transverse diffusion, particularly in the stratum corneum, was strongly dependent on the patch size and did not play a critical role in conventional patches.

## Nomenclature

## Symbols

A: age [a]
A_pt_: active area of the patch [m^2^]
c_i_^α^: drug concentration of substance α in material i [kg m^-3^]
c_sc,max_^α^: maximal concentration in the stratum corneum [kg m^-3^]
*c*_*pt,ini*_^*α*^: initial concentration in the patch [kg m^-3^]
d_sc_: thickness of stratum corneum [m]
d_ep_: thickness of epidermis [m]
d_vep_: thickness of viable epidermis [m]
d_pt_: thickness of patch [m]
D_i_^α^: diffusion coefficient/diffusivity of substance α in material i [m^2^ s^-1^]
G_bl,up_(t): uptake flow rate in blood at a specific point in time [kg s^-1^]
G_pt,rel_(t): release flow rate of patch at a specific point in time [kg s^-1^]
g_bl,up_(t): uptake flux across the skin into the blood at a specific point in time [kg m^-2^ s^-1^]
K_A/B_^α^: partition coefficient between material A and B for substance α
K_o/w_^α^: partition coefficient between octanol and water for substance α
K_i_^α^: drug capacity of substance α in material i [-]
L_pt_: length (or width) of patch (reservoir) [m]
L_sk_: length (or width) of skin [m]
m_pt,ini_: initial amount of drugs contained in the patch [kg]
m_pt,res_(t): remaining (residual) amount of drugs contained in the patch at a specific point in time [kg]
m_ep,stor_(t): total amount of drugs stored in the epidermis at a specific point in time [kg]
m_pt,rel_(t): cumulative amount of drugs released by the patch at a specific point in time [kg]
m_bl,up_(t): cumulative amount of drugs taken up by the blood flow at a specific point in time [kg] R diffusive resistance of a material [s m^-1^]
S_s_^α^: volumetric source term for substance α [kg m^-3^s^-1^]
S_U,Xj_: relative sensitivity of U to a change in X_j_
t: time [s]
t_1/2_: half-uptake-time [s]
U: process quantity
X_j_: model input parameter
Y_bl,up_: fractional drug release of the patch [-]

## Greek symbols

α: substance indicator
ψ^α^: drug potential of substance α [kg m^-3^]

## Subscripts

bl: blood
i: material indicator
ini: initial
sc: stratum corneum
ep: epidermis
vep: viable epidermis
ini: initial
fin: final
up: uptake
rel: release
stor: stored
sk: skin
pt: patch

## Abbreviations

HUT: half-uptake-time
TDDS: transdermal drug delivery systems

## Acknowledgments

This work was supported by the Novartis Research Foundation [grant “Virtual twinning for intelligent, personalized transdermal drug delivery”].

## AUTHOR CONTRIBUTIONS

T.D. and R.R. conceptualized the study and acquired funding; T.D. did the project administration; T.D. performed the investigation, developed the methodology, performed the validation, and executed the simulations with key input from L.D., who did exploratory simulation work during her MSc thesis; T.D. and A.T. supervised L.D. and F.B.; T.D. wrote the original draft of the paper and did the visualization, with key input from R.I.M. and F.B.; R.R., F.B., L.D., R.I.M. and A.T. performed critical review and editing.

## SUPPLEMENTARY MATERIAL

### 1 Derivation of conservation equation

The conservation equation was reformulated to another dependent variable to avoid numerical stability issues during calculation and to obtain a single equation that could be solved throughout the entire computational domain. In most studies (Table 1), in each material i (stratum corneum - sc, viable epidermis - ep, drug patch/reservoir - pt), the following conservation equations are solved for substance α:

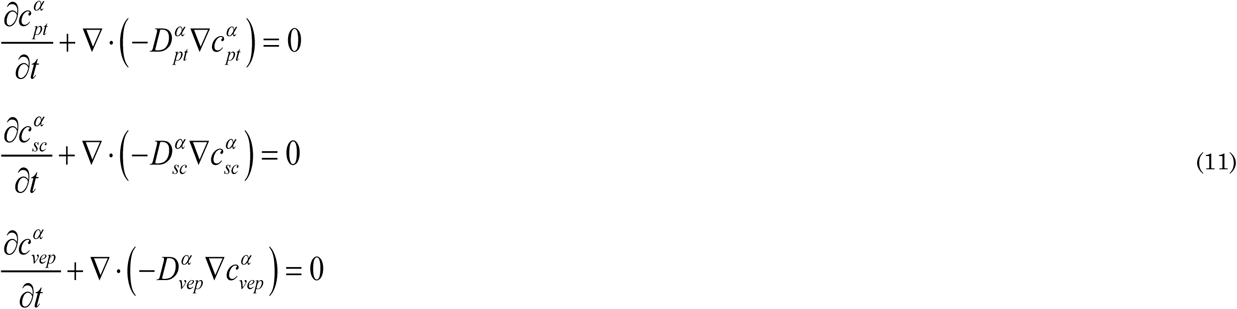

The concentrations in the different materials are linked via the partition coefficients by:

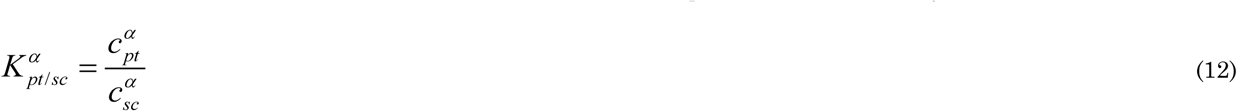

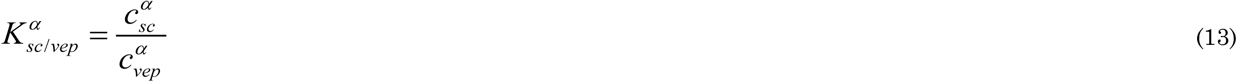

Additionally, flux continuity across the interfaces of different materials is assumed:

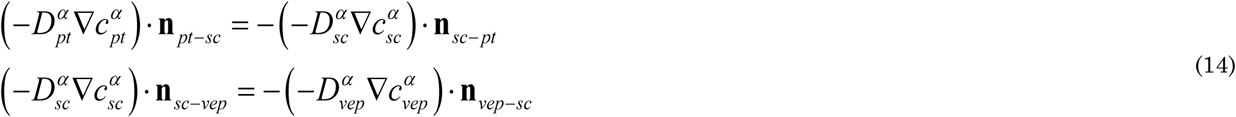

In the conservation equations above, we substituted the dependent variable drug concertation 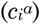 with the dependent variable potential 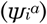 in each of these materials, based on following relations:

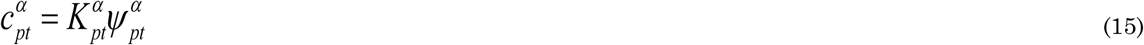

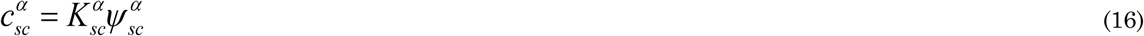

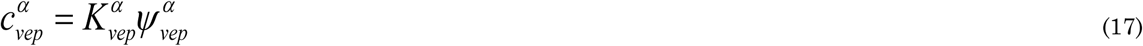

where 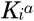 is termed the drug capacity in the material *i* [-]. This capacity is defined to be related to the partition coefficient in the following way:

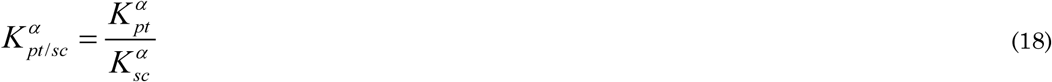

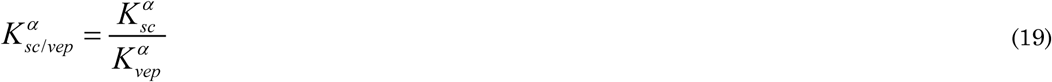

As such, when combining these equations, it follows that at the material interface:

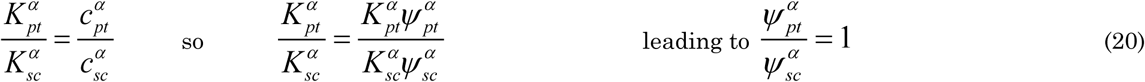

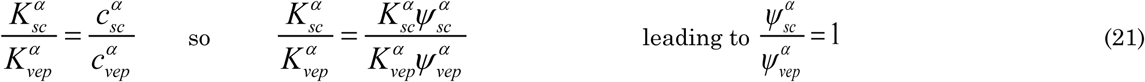

Thus, we get 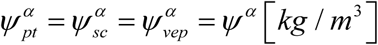. Hence, only one dependent variable (*ψ*^*α*^) needs to be solved for the entire domain, instead of several concentrations 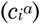 in each material *i* separately. When substituting Eq. (15), (16), and (17) in Eq. (1), we get for each material i:

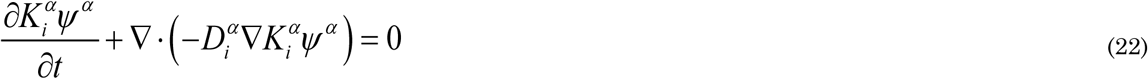

If we assume that 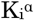 is constant within each material (so 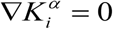, except at the border between materials), we can write:

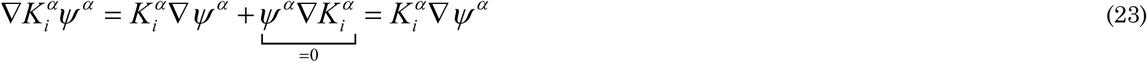

If we also assume that 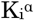 is constant over time, Eq. (22) becomes:

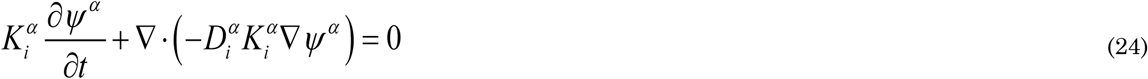

This equation is valid over the entire computational domain, with one dependent variable (*ψ*^*α*^), which is continuous throughout all materials and over all interfaces. Only 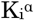 and 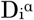 are material-specific, by which no explicit continuity of fluxes needs to be imposed.

### 2 Validation simulations

The drug release and uptake model was validated with previous experimental data [21]. This process was performed by quantifying how accurately the drug uptake kinetics were predicted with our mechanistic model. The experiment considered fentanyl uptake of a cylindrical drug reservoir (radius 9 mm, thickness 50.8 µm) through a cylindrical skin sample (human cadaver epidermis, radius 9.25 mm, thickness 50.8 µm). Both components were fitted into a Franz diffusion cell and monitored at 33°C for 72 h. Several initial drug concentrations in the patch were evaluated. For the validation simulations, 60 and 80 kg m^-3^ were considered. The flux in the experiments was determined over a specified time period of multiple hours by removing an aliquot of the receptor medium and analyzing the concentration via high-performance liquid chromatography (HPLC).

The conservation equations, boundary conditions, and initial conditions that were solved are similar to those detailed in section 2.1, but what follows is a brief description of the differences. A 3D cylindrical model of the setup was made. However, due to the very similar radius of the patch and the skin sample, the transport processes were quasi-one-dimensional. The small differences when comparing the 3D and 1D models for the validation study confirmed this fact (Figure 2). Thus, only the results for the 1D model are shown. The geometrical specifications and transport properties used in the simulations are indicated in Table 3. Note that the drug fluxes leaving the epidermis in experiments and simulations were scaled to the surface area at the patch-skin interface (surface 2, Figure 3), and not to that of the skin (which is slightly larger; surface 1).

### 3 Modeling assumptions

The impact of several modeling assumptions was evaluated. First, we evaluated whether a unidirectional (1D) transport model could be used as a simplification of the 3D transport problem. For this purpose, the base case for a 1D model (L_pt_ = L_sk_) was compared with the 3D model for the base case (L_sk_ = L_pt_ + 20 x (d_sc_ + d_vep_)). This comparison implies quantifying whether the transport in the transverse direction impacts the solution for a standard patch size. A comparison of the uptake flow (G_bl,up_(t)), the total amount that is taken up by the blood flow (m_bl,up_(t)), and the maximal concentration in the skin (c_sc,max_(t)), is shown in Figure S1, by quantifying the differences between both model types. The differences dropped below 1% within the first hour. This finding is not surprizing due to the large size of the patch: the ratio of the patch width to the epidermal thickness (stratum corneum and viable epidermis) was 471. For such very wide computational domains, a 1D approximation is clearly sufficiently accurate, because the edge effects do not affect the total released amount. Thus, in this study, the use of a 1D unidirectional transport model was justified. For smaller patch sizes, or in case the dermis would be included in the model, the use of 3D modeling should be re-evaluated, but this evaluation should be assessed on a case-by-case basis.

With regards to these findings, the impact of drug flow anisotropy in the stratum corneum was evaluated, because the diffusion coefficient in the transverse direction is much higher than the one in the longitudinal direction (Table 3). To this end, an isotropic versus anisotropic diffusion coefficient for the stratum corneum in the transverse direction was evaluated for a standard patch size. The base case (3D model, anisotropic diffusion coefficients) was compared with the case with isotropic diffusion coefficients. Here, the transverse diffusion coefficient was set equal to the longitudinal one, so much smaller than with the base case, namely 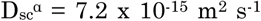. The differences in uptake flow rate (G_bl,up_(t)) were < 1%. Due to the large patch width, differences in transport in the transverse direction, which occur mainly at the patch edges, did not impact the total uptake flow rate.

Finally, the material properties used for the validation simulations (see [21]) differed from those used in the other simulations, which were taken from another study [26]. In the validation simulations, only one homogeneous epidermis layer was considered: the stratum corneum and the viable epidermis were embedded, which lumped their effect in one single diffusion and partition coefficient (Table 3). In our other simulations, the stratum corneum and the viable epidermis were discretely modeled, each with their diffusion and partition coefficient. We evaluated whether the utilized parameter set for the base case provided transport kinetics that were in the same range as the validated model parameter set. There was only a 20% difference in m_bl,up_(t) and G_bl,up_(t) between these two sources for material properties after the first 15 h. The difference in the total drug amount taken up at the end of the simulation was only 10%. Thus, both sets of material properties lead to similar transport kinetics. The main reason for this result is that the resistance to diffusion of both skin types for these two cases is very similar. This resistance R [s m^-1^] for a skin built out of *n* layers can be defined as:

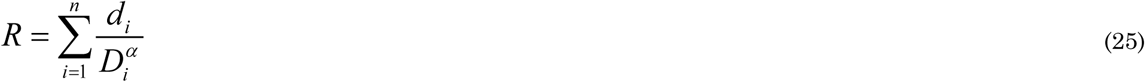

where *d*_*i*_ is the thickness of each layer [m]. The corresponding resistances for the diffusion coefficients of validation case and the base case were 0.733 x 10^9^ and 2.084 x 10^9^ s m^-1^, respectively. Due to these similar diffusional resistances, the validation case can be considered representative of the other simulations. The partition coefficients also differ, by which the amount of drug stored in the skin will be different for both cases: the largest amount is stored for the material properties taken from the validation case.

**Figure S1.**
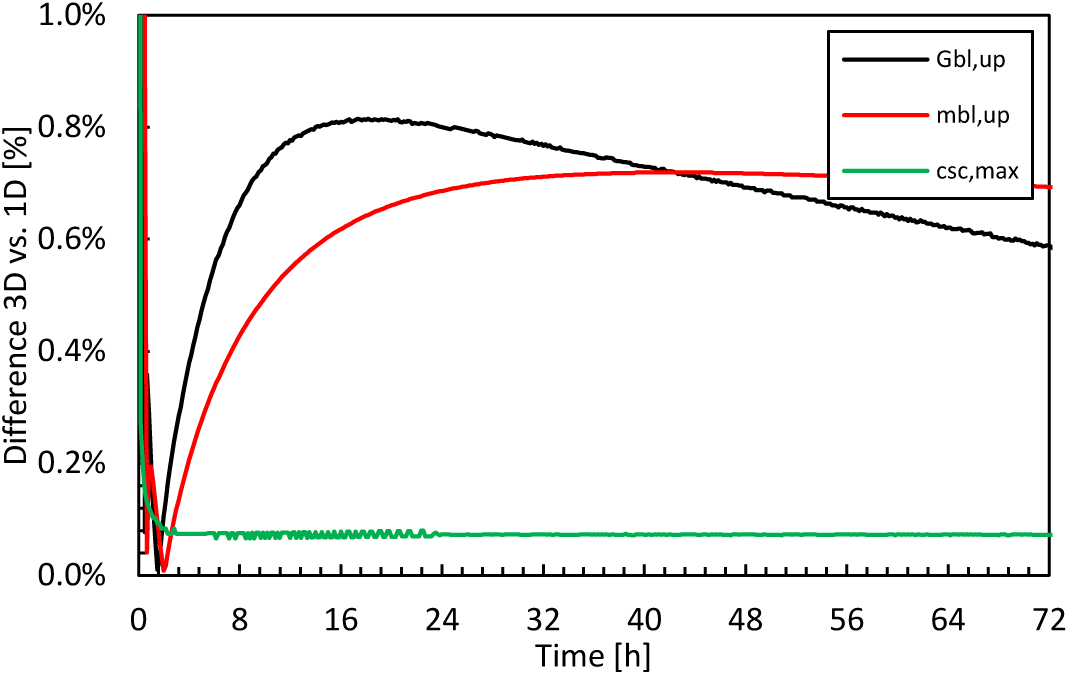
Differences of 3D and 1D models concerning the uptake flow (G_bl,up_), the total amount taken up by the blood flow (m_bl,up_) and the maximal concentration in the skin (c_sc,max_) as a function of time.

### 4 Impact of the dermis layer

The impact of including the diffusion in the dermis on the diffusive drug uptake is illustrated by evaluating different dermis thicknesses, namely 100, 200, 400, 800, 1,600 and 3,200 μm. Only diffusion (and not blood flow) was modeled in the dermis. The uptake flux (g_bl,up_) and the amount taken up by the blood flow (m_bl,up_) are shown in Figure S2. Since no blood flow in the dermis was modeled, the time lag before the drug reaches the blood, i.e., the lower boundary of the computational domain increased significantly with increasing dermis thickness (Figure S2a). Furthermore, the drug amount taken up over 72 h decreased with increasing dermis thickness. Correspondingly, the stored drug amount increased (Figure S2b) and the peak in the uptake flux decreased.

From these results, the dermal blood flow should be included in the model when the dermis is explicitly modeled because the differences found in Figure S2a for the same drug reservoir were very large. It is not realistic that, if the size of the computational domain is increased, the predicted drug uptake by the patient decreases to such an extent. Hence, modeling the dermis without drug extraction by the blood flow likely underpredicts the drug uptake rate. Due to the large thickness and volume of the dermis, only modeling diffusive transport can also overpredict the stored drug amount and extensive transversal drug spreading in the dermis.

**Figure S2.**
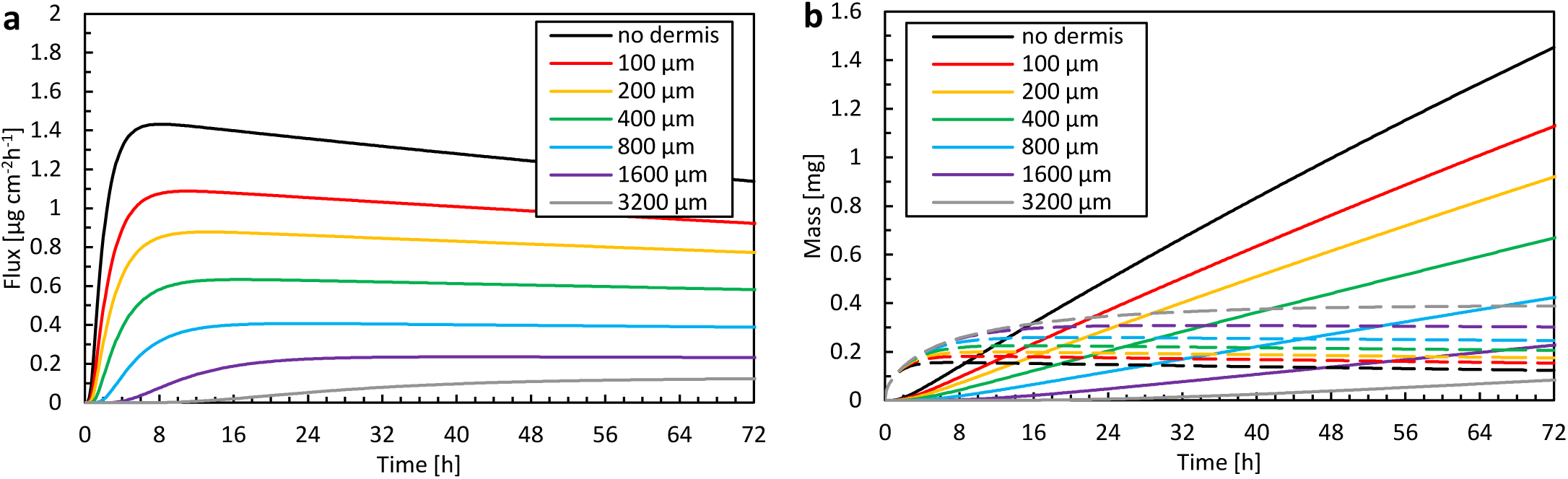
The total amount that is taken up by uptake flux (g_bl,up_) and the blood flow (m_bl,up_) as a function of time for different thicknesses of the dermis.

